# eNOS-induced vascular barrier disruption in retinopathy by c-Src activation and tyrosine phosphorylation of VE-cadherin

**DOI:** 10.1101/2020.11.24.396077

**Authors:** Takeshi Ninchoji, Dominic T. Love, Ross O. Smith, Marie Hedlund, Dietmar Vestweber, William C. Sessa, Lena Claesson-Welsh

**Affiliations:** Uppsala University, Rudbeck Laboratory, Department of Immunology, Genetics and Pathology, Uppsala, Sweden; Max Planck Institute for Molecular Biomedicine, Münster, Germany; Yale University School of Medicine, Department of Pharmacology and Vascular Biology and Therapeutics Program, New Haven, USA

**Keywords:** Medicine, Cell Biology

## Abstract

Hypoxia and the production of vascular endothelial growth factor A (VEGFA) promote blood vessel leakiness and edema in ocular diseases. Therapeutics targeting VEGFA suppress leakiness and edema but aggravate hypoxia; therefore, new therapeutics are needed. We examined the role of endothelial nitric oxide synthase (eNOS) in pathological neovascularization and vessel permeability during oxygen-induced retinopathy. NO formation was suppressed chemically using L-NMMA, or genetically, in eNOS serine to alanine (S1176A) mutant mice, resulting in reduced retinal neoangiogenesis. Both strategies resulted in reduced vascular leakage by stabilizing endothelial adherens junctions through suppressed phosphorylation of vascular endothelial (VE)-cadherin Y685 in a c-Src-dependent manner. Intervention treatment by a single dose of L-NMMA in established retinopathy restored the vascular barrier and prevented leakage. We conclude that eNOS induces destabilization of adherens junctions and vascular hyperpermeability by converging with the VEGFA/VEGFR2/c-Src/VE-cadherin pathway and that this pathway can be selectively inhibited by blocking NO formation.

## Introduction

Pathological neovascularization is intimately associated with the progression of several retinal diseases, including retinopathy of prematurity, diabetic retinopathy, and exudative age-related macular degeneration. Neovascularization occurs in response to hypoxia, tissue ischemia and the consequent production of angiogenic agonists, such as vascular endothelial growth factor A (VEGFA), a potent inducer of vessel formation and vascular leakage (Campochiaro, 2015; Semenza, 2012). The new vessels formed during retinal ischemia are often dysfunctional and fail to stabilize (Fruttiger, 2007; Krock, Skuli, & Simon, 2011), leading to vessel leakage or hemorrhaging, and to retinal detachment, visual impairment and even blindness. Therefore, suppression of neoangiogenesis and thereby, retinal edema, is a therapeutic goal in the treatment of ischemic eye diseases (Daruich et al., 2018).

A number of therapeutic options designed to neutralize VEGFA by preventing binding to its receptor, VEGF receptor 2 (VEGFR2), such as bevacizumab, ranibizumab, and aflibercept, decrease neovascular formation as well as edema (Mintz-Hittner, Kennedy, Chuang, & Group, 2011). However, they do not correct the underlying hypoxia, in fact, vessel regression may instead further aggravate hypoxia. Moreover, in many cases anti-VEGF therapies can induce an elevation in intraocular pressure and hemorrhaging (Wells et al., 2015). The repeated intravitreal injections of anti VEGF-therapy present a potential for infections and scarring (Patel, Cholkar, Agrahari, & Mitra, 2013), in addition, side effects including disrupted neural development in infants have been reported (Morin et al., 2016). Thus, even though the current therapy improves vision for many patients in the early phases of disease, there is a clear need for developing new treatments to suppress proangiogenic stimuli while also achieving a long-lasting effect with safe administration.

Angiogenesis and vascular permeability in the retina are initiated primarily by the VEGFA/VEGFR2 signaling pathway. VEGFR2 is found on blood vascular endothelial cells but also on neuronal cells in the retina, explaining the side effect of VEGFA/VEGFR2 suppression on neural development in premature infants. VEGFR2 activity is initiated through VEGFA-induced receptor dimerization, kinase activation, phosphorylation of tyrosine residues in the receptor intracellular domain and activation of signaling pathways. The VEGFR2 phosphotyrosine-initiated signaling pathways are now being unraveled. Thus, the phosphorylation of Y1212 in VEGFR2 is required for activation of phosphatidyl inositol 3 kinase and AKT (Testini et al., 2019), while phosphorylation of Y949 is required for activation of the c-Src pathway and regulation of vascular permeability through vascular endothelial (VE)-cadherin (Li et al., 2016). VE-cadherin is the main component of endothelial adherens junctions, strongly implicated in regulation of vascular permeability, leakage and associated edema (Giannotta, Trani, & Dejana, 2013). In particular, phosphorylation of the Y685 residue in VE-cadherin correlates with VEGFA-induced vascular hyperpermeability (Smith et al., 2020; Wessel et al., 2014), where Y685 phosphorylation leads to the dissociation of the homophilic interactions between VE-cadherin molecules expressed on adjacent endothelial cells (Giannotta et al., 2013).

Endothelial nitric oxide synthase (eNOS), activated downstream of VEGFA through phosphorylation on S1177 (S1176 in mice) by AKT (Fulton et al., 1999), produces nitric oxide (NO) and regulates vascular permeability. Both mice with a constitutive eNOS gene (*Nos3*) inactivation (Fukumura et al., 2001), and mice expressing an eNOS serine to alanine point mutation (S1176A) (*Nos3*^*S1176A/S1176A*^) (Di Lorenzo et al., 2013) show reduced extravasation of bovine serum albumin in the healthy skin in response to VEGFA challenge. An important mediator of the effect of eNOS-generated NO is the relaxation of perivascular smooth muscle cells, leading to increased vessel diameter and enhanced blood flow and thereby flow-driven vascular sieving (Sessa, 2004). In addition, eNOS activity correlates with tyrosine phosphorylation of VE-cadherin in cultured endothelial cells (Di Lorenzo et al., 2013), providing a mechanism for how eNOS activity may directly affect vascular permeability, distinct from vasodilation.

In contrast to the skin vasculature, the healthy retinal vasculature is protected by a stringent blood-retinal barrier. In retinal diseases, the barrier is disrupted, leading to increased vascular permeability (Zhao, Nelson, Betsholtz, & Zlokovic, 2015). In accordance, NO and related reactive oxygen species (ROS) are important pathogenic agents in retinopathy (Opatrilova et al., 2018). However, the molecular mechanisms whereby eNOS/NO interferes with the retinal vascular barrier and contributes to pathological vascular permeability in eye disease have remained unexplored. Here, we show that suppressed NO formation via the use of the competitive NOS inhibitor, L-NMMA, or an eNOS mutant, S1176A, negates neovascular tuft formation and vascular leakage during retinal disease. Mechanistically, NO promotes c-Src Y418 phosphorylation at endothelial junctions and phosphorylation of VE-cadherin at Y685, required for dismantling of adherens junctions. Inhibition of NO formation by L-NMMA treatment suppresses vascular leakage also from established neovascular tufts, separating regulation of leakage from the angiogenic process as such. Mice expressing a VE-cadherin tyrosine to phenylalanine mutation (VEC-Y685F) are resistant to eNOS inhibition, in support of that NO regulates adherens junctions through direct effects on c-Src and VE-cadherin. These data suggest that eNOS/NO promote vascular permeability not only through the established effect on vascular smooth muscle relaxation and increased flow-driven permeability to solute and small molecules in the precapillary arterial bed, but also through disruption of adherens junctions allowing leakage of larger molecules from the postcapillary venular bed.

## Results

### Reduced neoangiogenic tuft formation in C57BL/6 OIR with suppressed NO formation

To determine how eNOS inhibition affects pathological angiogenesis in retinopathy in C57BL/6J mice, pups were exposed to 75% oxygen from P7-P12 (hyperoxic period) where after they were placed at normal, atmospheric conditions (21% oxygen; relative hypoxic period) from P12-P17 (Figure 1A). During the P7-P12 hyperoxic period, VEGFA expression is suppressed, leading to endothelial cell death and avascularity in the superficial vessel layer (reviewed in (Scott & Fruttiger, 2010)). The relative decrease in oxygen concentration upon return to normal atmosphere at P12-P17 induces hypoxia inducible factor-dependent gene regulation, causing oxygen-induced retinopathy (OIR) and the formation of neoangiogenic tufts (L. E. Smith et al., 1994) (see Figure 1B for schematic outline).

**Figure 1.**
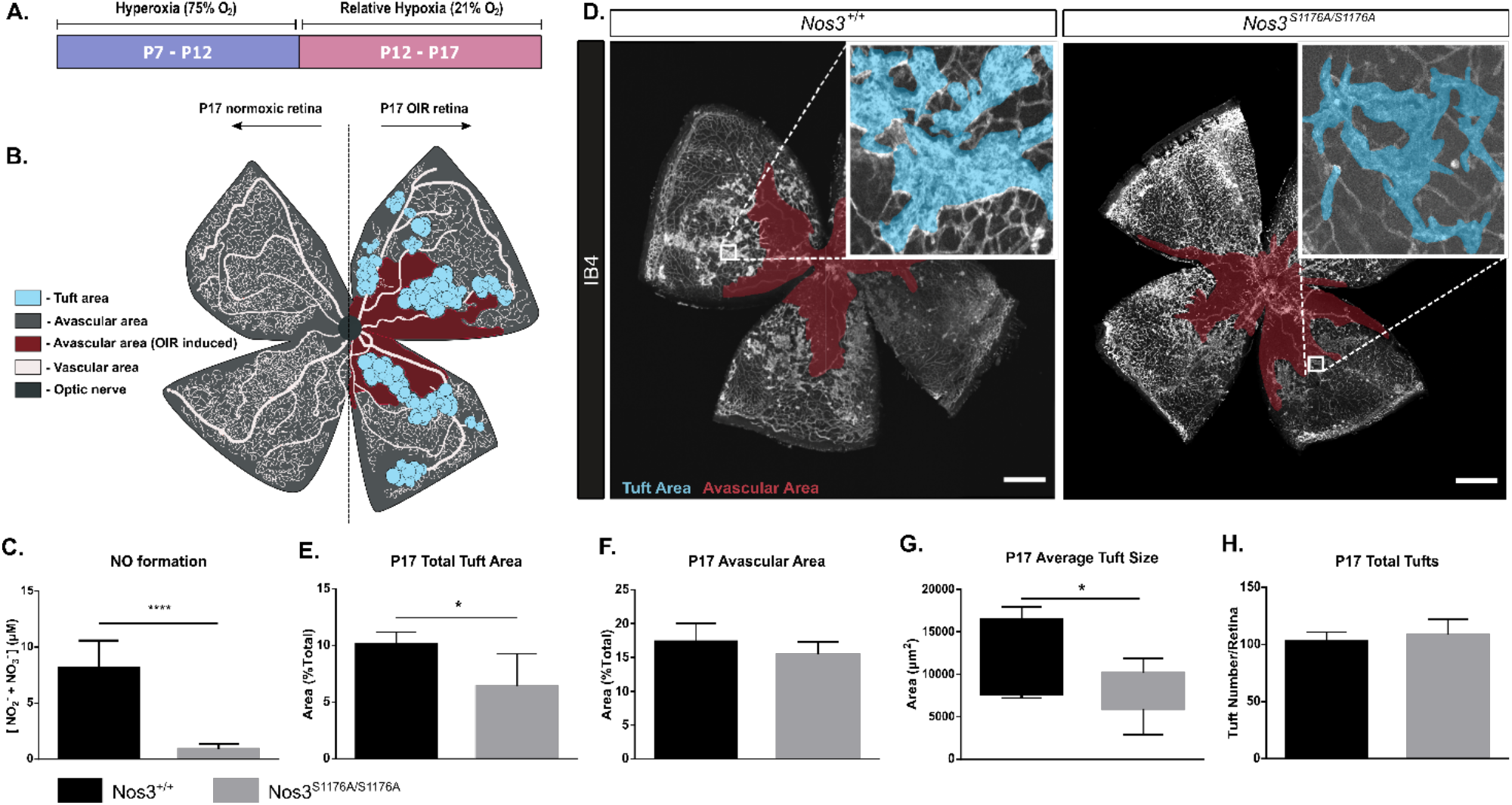
Suppressed tuft formation in the Nos3S1176A/S1176A retina after OIR-challenge. A. Outline of OIR-challenge protocol; pups were placed in 75% O2 (hyperoxia) between P7-P12, followed by return to normal atmosphere (relative hypoxia) until P17. B. Schematic representation of vascular abnormalities after OIR in P17 retinas. C. Nitric oxide formation determined using a Griess assay, expressed as the combined concentration of nitrite and nitrate, the end-products of NO, reacting with molecules in biological fluids. Mean ±S.E.M. n = 3 mice/genotype. **** = p <0.0001; t-test. D. Representative images of whole mount retinas from Nos3^+/+^ and Nos3^S1176A/S1176A^ mice, collected at P17 after OIR-challenge, stained with isolectin B4 (IB4). Avascular area is marked in magenta and tufts in blue. Scale bar = 500 μm. E-F.Tuft area (E) and avascular area (F) expressed as percentage of total vascular area at P17. G,H. Tuft size in μm2 (G) and total number/FOV in P17 mice (H). For E-H: mean ±S.E.M. n = 7 (Nos3^+/+^) and 5 (Nos3^S1176A/S1176A^) mice. * = p < 0.05; t-test. Figure 1 – source data 1. Raw data on retina vascular parameters and body weights from OIR experiments on *Nos*^*+/+*^ and *Nos3*^*S1176A/S1176A*^ mice Figure 1 – supplement figure 1. Postnatal development of *Nos3*^*+/+*^ and *Nos3*^*S1176A/S1176A*^ retinal vasculature Figure 1 – supplement figure 2. Retina development in *Nos3*^*+/+*^ and *Nos3*^*S1176A/S1176A*^ P12 pups Figure 1 – supplement figure 3. Expression of Nos2, Nos3 and Vegfa in *Nos3*^*+/+*^ and *Nos3S1176A/S1176A* retinas

To specifically address the role of eNOS in vascular retinal disease, we used a genetic model in which eNOS S1176 (mouse numbering (Fulton et al., 1999); S1177 in human) is replaced by alanine (A) (Schleicher et al., 2009). Phosphorylation of eNOS on this serine residue is a prerequisite for eNOS-driven NO production which was verified using a Griess assay on isolated endothelial cells from Nos3^+/+^ and *Nos3*^*S1176A/S1176A*^ mice (Fig. 1C). Mice were subjected to the OIR regimen (Fig. 1D). After OIR-challenge, *Nos3*^*S1176A/S1176A*^ P17 retinas showed reduced pathological tuft area compared to Nos3^+/+^ (Figure 1E), while the extent of avascularity was the same (Figure 1F). The average size of individual tufts was reduced in the *Nos3*^*S1176A/S1176A*^ pups (Figure 1G) while the total number of tufts formed after OIR was unaffected (Figure 1H). See Figure 1 – source data 1, for vascular parameters and body weights of mice.

The suppressed formation of neoangiogenic tufts in the absence of eNOS S1176 phosphorylation was not due to developmental defects as *Nos3*^*S1176A/S1176A*^ mice showed normal postnatal vascular development. Vascular plexus area and outgrowth in the retina, tip cell number and appearance, as well as branch points were all similar between the wild type and the *Nos3*^*S1176A/S1176A*^ retinas (Figure 1 – figure supplement 1A-G). At P12 after OIR-challenge, there was also no difference in the degree of avascularity in the retina between *Nos3*^*S1176A/S1176A*^ and Nos3^+/+^ pups, indicating that the strains responded similarly to the hyperoxic challenge (Figure 1 – figure supplement 2A, B). Importantly, the reduced tuft area in the *Nos3*^*S1176A/S1176A*^ condition was not a result of differences in *Nos3* or *Nos2* expression between the *Nos3*^+/+^ and *Nos3*^*S1176A/S1176A*^ mice before or after the OIR-challenge (Figure 1 – figure supplement 3A, B). Reduced tuft area was also not a result of reduced VEGFA-production as an equally induced level of VEGFA was seen in the mutant and wildtype mice (Figure 1 – figure supplement 3C). It should also be noted that the low relative expression level of *Nos2* (encoding inducible nitric oxide synthase (iNOS)) compared to *Nos3* (Figure 1 – figure supplement 3D, E) emphasizes the primary role of eNOS as a source of endothelial NO, both in the unchallenged and OIR-treated condition.

### VEGFA induces eNOS phosphorylation and activity

VEGFA produced during the relative hypoxia phase (P12-P17) is an important instigator of edema in retinopathy (Connor et al., 2009; Dor, Porat, & Keshet, 2001), which is characterized by leaky and dysfunctional vessels. VEGFA-mediated AKT activation leads to phosphorylation of eNOS at S1177 (Chen & Meyrick, 2004; Schleicher et al., 2009). In agreement, eNOS was phosphorylated on S1177 in VEGFA-treated human retinal microvascular endothelial cells (HRMEC). Induction of eNOS phosphorylation appeared with similar kinetics but slightly more potently by VEGFA than by the inflammatory mediator histamine (Figure 2 – figure supplement 1A, B; see Figure 2 – figure supplement 1C, D for antibody validation), a well-known inducer of eNOS activity (Thors, Halldorsson, & Thorgeirsson, 2004). eNOS phosphorylation was accompanied by NO production in response to VEGFA stimulation, as assessed using the fluorescent probe, DAF-FM diacetate added to the HRMEC culture medium. NO accumulated significantly by 1 min stimulation and still persisted at 10 min (Figure 2 – figure supplement 1E). DAF-FM fluorescence, and therefore NO production, was blocked by incubating cells with L-NMMA (Figure 2 – figure supplement 1F), to the level of the untreated control. Combined, these data show that VEGFA is a potent inducer of eNOS activity and NO production.

**Figure 2.**
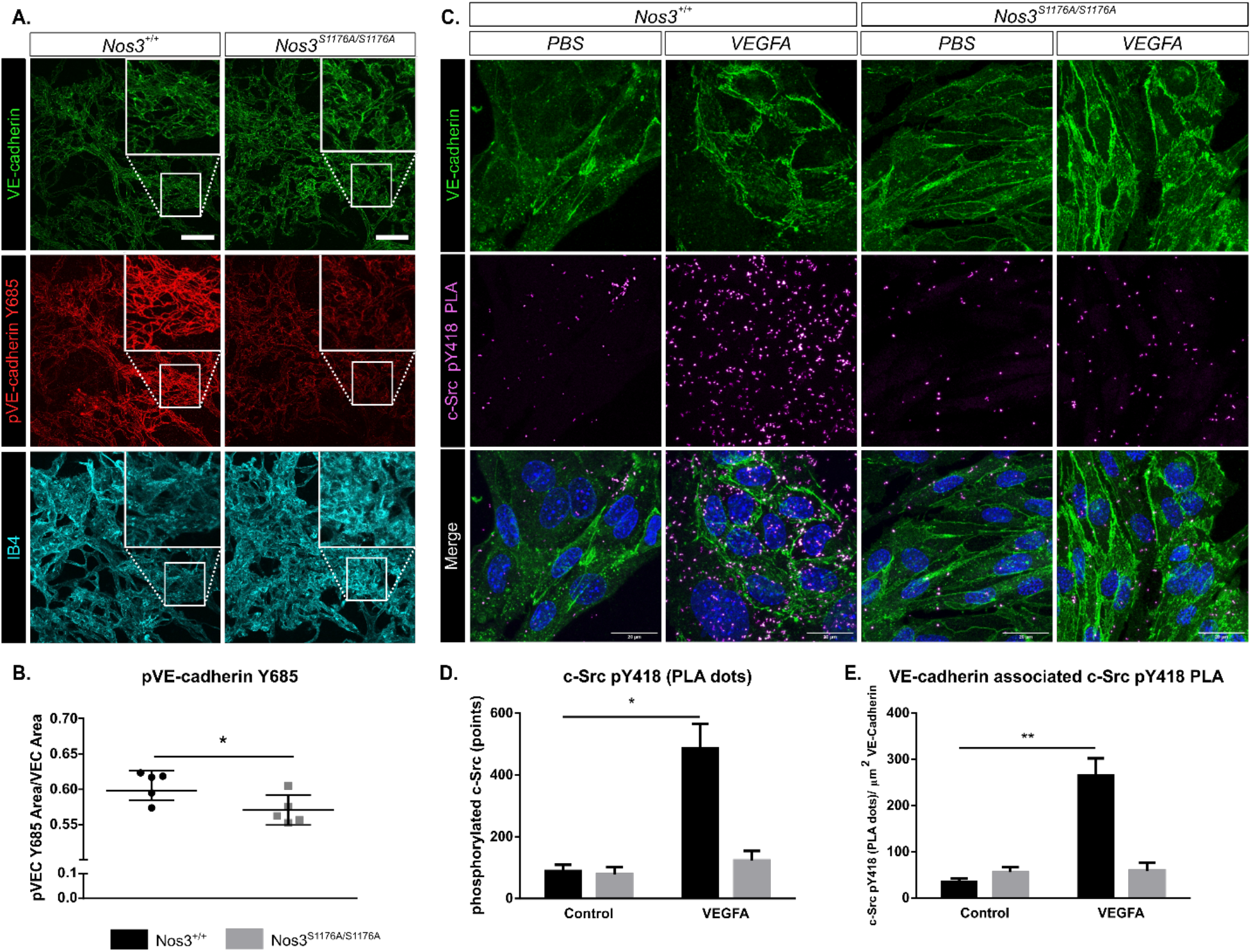
Suppressed c-Src Y418 and VE-cadherin Y685 phosphorylation in *Nos3S1176A/S1176A* retinas. A. Representative maximum intensity projections of tufts from *Nos3*^*+/+*^ and *Nos3*^*S1176A/S1176A*^ retinas immunostained for VE-cadherin (green), pY685 VE-cadherin (red) and isolectin B4 (IB4; cyan). Scale bar = 50 μm. B. Ratio of pY685 positive area/total VE-cadherin positive area. Mean ±S.E.M n = 3-6 images per group from 4 (*Nos3*^*+/+*^) and 6 (*Nos3*^*S1176A/S1176A*^) mice, 3 independent experiments, * = p < 0.05; t-test. C. Representative images of VE-cadherin staining (green) and proximity ligation assay (PLA) to detect c-Src pY418 (magenta) in isolated mouse lung endothelial cells (iEC) from *Nos3*^*+/+*^ and *Nos3*^*S1176A/S1176A*^ mice. Scale bar = 20 μM. D. c-Src pY418 PLA spots detected in PBS and VEGFA (100 ng/mL)-treated iECs from *Nos3+/+* and *Nos3*^*S1176A/S1176A*^ mouse lungs. E. c-Src pY418 PLA spots co-localized with VE-cadherin (green), normalized against total VE-cadherin area in the field of view. Mean ±S.E.M. n = 4 (*Nos3*^*+/+*^) and 4 (*Nos3*^*S1176A/S1176A*^) mice, from 3 separate experiments. * = p < 0.05; two-way ANOVA, Sidak’s multiple comparisons test. Figure 2 – supplement figure 1. VEGFA induced eNOS phosphorylation and activity *in vitro* Figure 2 – supplement figure 2. c-Src pY418 immunostaining and PLA controls

### VE-cadherin phosphorylation at Y685 is reduced in *Nos3*^*S1176A/S1176A*^ vessels after OIR due to the inhibition of c-Src Y418 phosphorylation

VEGFA/VEGFR2 signaling and vessel leakage correlates with phosphorylation of VE-cadherin on Y685 (Orsenigo et al., 2012; Smith et al., 2020; Wessel et al., 2014). The level of pY685 VE-cadherin was examined by immunostaining of Nos3^+/+^ and *Nos3*^*S1176A/S1176A*^ retinas at P17 after OIR-challenge. pY685 VE-cadherin immunostaining, normalized to the total VE-cadherin area, was significantly lower in *Nos3*^*S1176A/S1176A*^ tufts than in the WT tufts (Figure 2A, B).

Phosphorylation of VE-cadherin on Y685 is dependent on the cytoplasmic tyrosine kinase c-Src (Wallez et al., 2007). The NO-generating reagent, SNAP, can increase the levels of activated c-Src phosphorylated on Y418 in fibroblast cultures (Rahman et al., 2010) indicating a potential role for NO in c-Src activation. We therefore tested whether eNOS 1176 phosphorylation correlates with phosphorylation of c-Src at Y418. However, immunostaining for c-Src pY418 failed to reveal differences in pY418 c-Src levels between Nos3^+/+^ and *Nos3*^*S1176A/S1176A*^ retinas (Figure 2 – figure supplement 2A, B). This was most likely due to that the pY418 antibody recognized several related Src-family molecules such as Yes and Fyn. Therefore, endothelial cells were isolated from lungs of Nos^+/+^ and *Nos3*^*S1176A/S1176A*^ mice and treated or not with VEGFA. To specifically determine induction of c-Src pY418, and not related Src family kinases, we employed the proximity ligation assay (PLA) (Soderberg et al., 2006), using oligonucleotide-ligated secondary antibodies detecting primary antibodies against murine c-Src protein and the conserved pY418 residue. The c-Src pY418 PLA was combined with counterstaining for VE-cadherin (Figure 2C). As shown in Figure 2D, VEGFA stimulation increased PLA spots, representing c-Src pY418, at least five-fold in the isolated endothelial cells (iECs) from Nos^+/+^ mice, but not in iECs from *Nos3*^*S1176A/S1176A*^ mice (see Figure 2 – figure supplement 2C for PLA controls). The c-Src pY418 PLA spots co-localized with VE-cadherin immunostaining (Figure 2E).

These data indicate that eNOS S1176 phosphorylation and its role in the formation of NO are essential for the accumulation of active c-Src at endothelial junctions to induce the phosphorylation of VE-cadherin at Y685.

### Suppressed vascular leakage in *Nos3*^*S1176A/S1176A*^ retinas after OIR-challenge

In the retina, the blood retina barrier (BRB) controls vascular permeability, however, the BRB is disrupted in retinopathies, causing edema and vision loss (Klaassen, Van Noorden, & Schlingemann, 2013; Zhao et al., 2015). Edema correlates with reduced vessel permeability, which is dependent on the phosphorylation status of VE-cadherin (Wessel et al., 2014) and c-Src activity (Wallez et al., 2007). To assess the role for eNOS specifically in vessel leakage after hypoxia-driven VEGFA production, 25 nm fluorescent microspheres were injected in the tail vein of P17 wildtype and *Nos3*^*S1176A/S1176A*^ mice, after OIR-challenge. Confocal image analysis showed accumulation of microspheres outside the vascular tufts, in agreement with enhanced vessel leakage upon OIR (Figure 3A). The accumulation of microspheres normalized to tuft area, was significantly lower in *Nos3*^*S1176A/S1176A*^ retinas compared to Nos3^+/+^ (Figure 3B; Figure 3 – source data 1).

**Figure 3.**
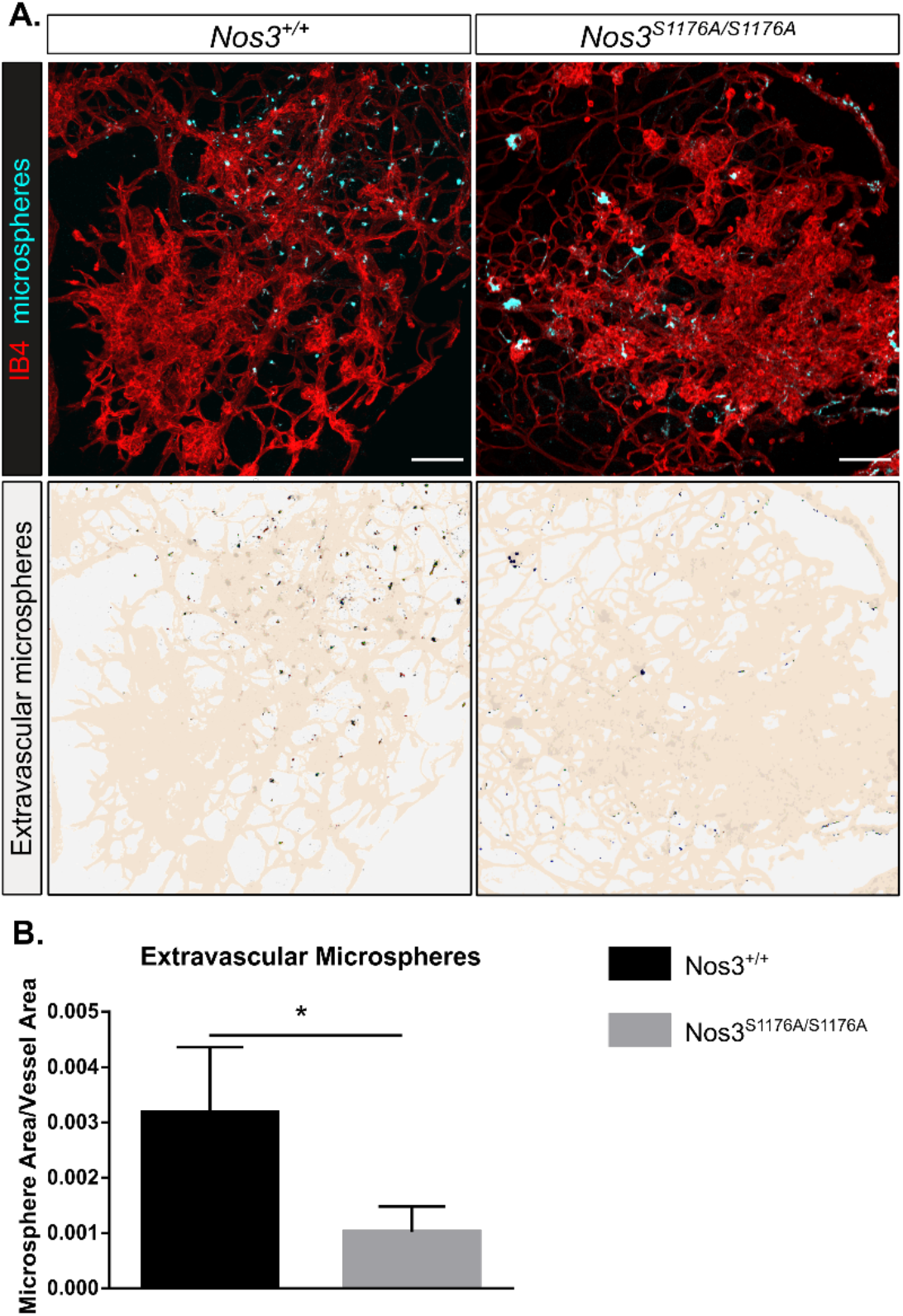
Suppressed microsphere leakage in *Nos3*^*S1176A/S1176A*^ retinas. A. Representative images of tufts from *Nos3*^*+/+*^ and *Nos3*^*S1176A/S1176A*^ mice immunostained for isolectin B4 (IB4; red), demonstrating leakage of tail-vein injected FITC-conjugated 25 nm microspheres (cyan) around the tufts. Scale bar = 100 μM. Lower panels show leakage maps. Heat mapped dots that do not overlap with vessels (beige) are considered extravascular. B. Quantification of the average area of extravascular microspheres normalized to IB4 area. Mean ±S.E.M. n = 4 (*Nos3*^*+/+*^) and 4 (*Nos3*^*S1176A/S1176A*^) mice, 6 – 15 images per mouse. * = p < 0.05, t-test. Figure 3 – source data 1. Raw data on retina vascular parameters and body weights from *Nos3*^*+/+*^ and *Nos3*^*S1176A/S1176A*^ mice injected with microspheres

### Reduced tuft area as a result of pharmacological inhibition of NO formation

The *Nos3*^*S1176A/S1176A*^ mouse is unable to produce NO in the endothelium due to the non-phosphorylatable alanine replacing S1176. As we could not unequivocally exclude that the vascular effects observed in the *Nos3*^*S1176A/S1176A*^ mutant were dependent on non-NO synthesis events linked to S1176 phosphorylation, we tested the effect of the cell permeable NOS inhibitor Nω-Methyl-L-arginine (L-NMMA), which inhibits NO formation from all three NOS variants (eNOS, inducible NOS and neuronal NOS).

Intraperitoneal injection of L-NMMA (20 μg/g body weight) were given daily during P12-17 (injections on days P12-P16), i.e. treatment was initiated before pathological neovessels were established (prevention therapy). L-NMMA treatment significantly reduced the area of vascular tufts formed by P17 (Figure 4A, B) (Figure 4 – source data 2) but did not affect the avascular area (Figure 4C). The average tuft size was decreased (Figure 4D) while the total number of individual tufts increased with L-NMMA treatment (Figure 4E). As smaller tufts can fuse to form larger structures (Prahst et al., 2020), the increase in individual tufts in the L-NMMA-treated litter mates may reflect the suppressed growth and fusion of the tufts. We conclude that while chemical eNOS inhibition suppressed the growth of tufts, it did not block formation of tufts *per se*.

**Figure 4.**
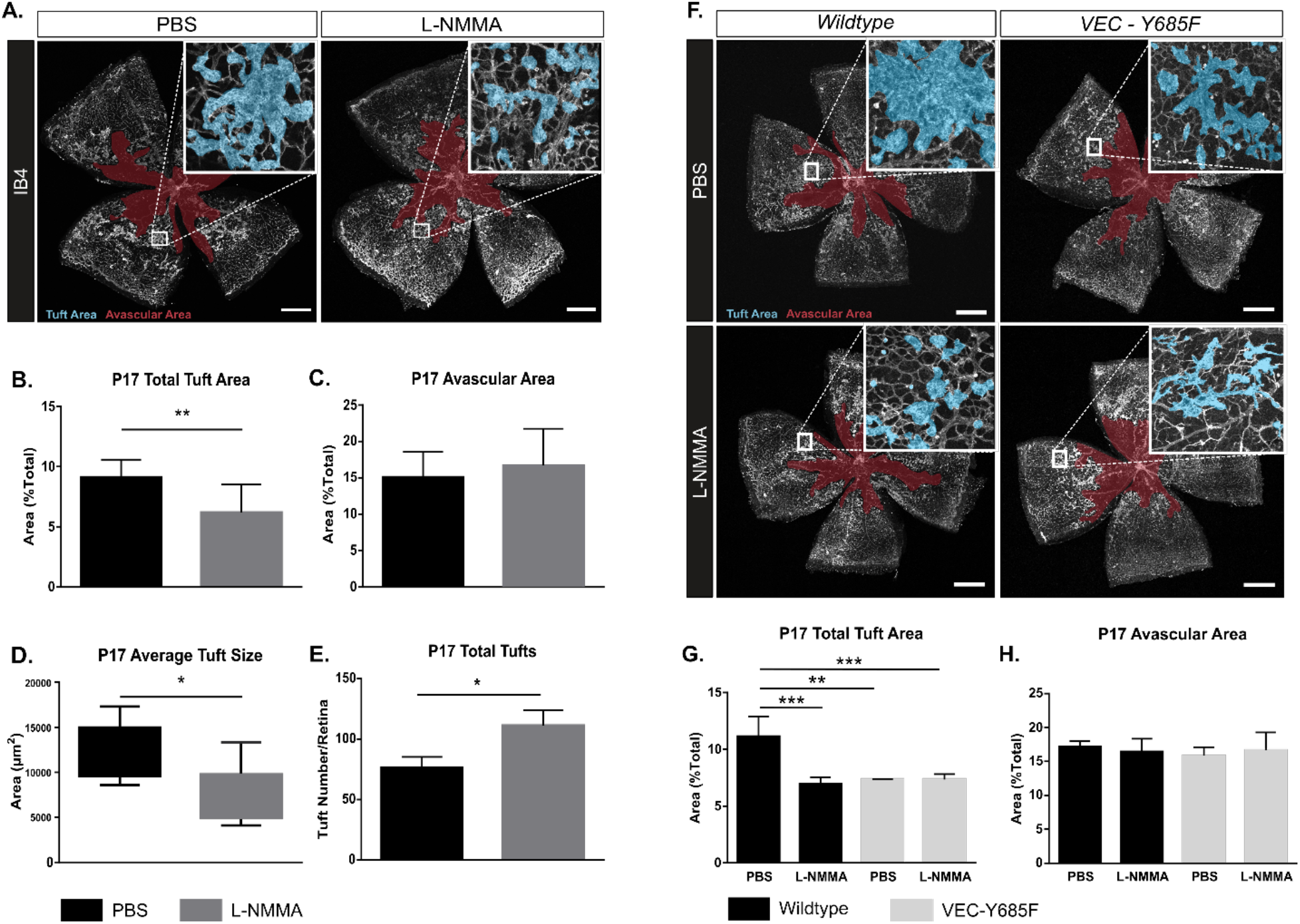
OIR-challenged mice treated with NO inhibitor L-NMMA. A. Representative images of whole mount retinas from PBS and L-NMMA treated (P12-P16) wildtype C57Bl/6 mice, collected on P17 after OIR challenge, and stained with isolectin B4 (IB4). Avascular area as a result of OIR is marked in magenta and tufts in blue. Scale bar = 500 μm. B, C. Tuft area and avascular area expressed as percentage of total vascular area at P17. D, E. Tuft size in μm^2^ and total number of tufts/field of vision at P17. Mean ±S.E.M. n = 8 (PBS) and 9 (L-NMMA) treated mice. *, ** = p < 0.05, 0.01; t-test. F. Representative images of whole mount retinas from OIR-challenged wildtype and VEC-Y685F mice injected with PBS or L-NMMA during P12-P16. Immunostaining for isolectin B4 (IB4) at P17. Avascular area is marked in magenta and tufts in blue. Scale bar = 500 μm. G. Tuft area normalized to total vascular area in PBS or L-NMMA-treated wildtype and VEC-Y685F retinas. H. Avascular area normalized to total vascular area. Mean ±S.E.M. n = 8 (VEC^+/+^) and 8 (VEC^Y685F/Y685F^) mice. **, *** = p < 0.01, 0.001; two-way ANOVA, Sidak’s multiple comparison test. Figure 4 – source data 1. Raw data on retina vascular parameters and body weights from OIR experiments on PBS and L-NMMA treated mice Figure 4 – source data 2. Raw data on retina vascular parameters and body weights from OIR experiments on VEC^+/+^ and VEC-Y685F mice

### NO and VE-cadherin Y685 phosphorylation operate on the same pathway regulating vascular leakage

To further explore the relationship between NO and VE-cadherin pY685 in the formation of leaky, pathological vessels, we used mice expressing mutant VE-cadherin wherein phosphorylation at position 685 is abolished by exchanging the tyrosine (Y) for phenylalanine (F), termed VEC-Y685F (Wessel et al., 2014). VEC-Y685F mice show suppressed induction of vascular leakage in the healthy skin (Wessel et al., 2014). We hypothesized that if NO modulates vascular leakage and tuft formation via a non-VE-cadherin Y685 pathway, L-NMMA would impart an additional reduction in tuft area to OIR-challenged VEC-Y685F mice. To test whether the VEC-Y685F mouse would respond to NOS inhibition, L-NMMA (20 μg/g body weight) was administered by intraperitoneal injection of wildtype and VEC-685F mice during the relative hypoxic period (injections on days P12-P16). Results show that L-NMMA treatment did not further suppress tuft formation in Y685F mice at P17. The reduction in tuft area was similar, about 50%, in PBS and L-NMMA-treated Y685F retinas and comparable to that seen in L-NMMA-treated wildtype mice (Figure 4F, G) (Figure 4 – source data 2). The avascular area remained unaffected by all treatments (Figure 4H).

### Single-dose L-NMMA decreases vascular leakage in the retina

We next aimed to mimic a clinical situation by administering L-NMMA to OIR-challenged wild type mice with established pathological vessels (intervention therapy). Mice were given one injection of L-NMMA (60 μg/g body weight) at P16. At P17, microspheres were injected and after 15 min, the experiment was terminated. The area of extravascular microspheres, assessed after normalization to tuft area (Figure 5A, B) or to total microsphere area (Figure 5A, C) was reduced by 50-60% after the single-dose treatment with L-NMMA compared to PBS. Of note, the total microsphere area was not affected by the L-NMMA treatment (Figure 5D), indicating that microspheres to a large extent were present in the vascular lumen in the L-NMMA treated condition while in the PBS control, they had crossed the disrupted barrier to the extravascular space. Importantly, the tuft area was not affected by the L-NMMA treatment (Figure 5E) (Figure 5 – source data 1). Thus, these data indicate that leakage could be suppressed even from established neovascular structures.

**Figure 5.**
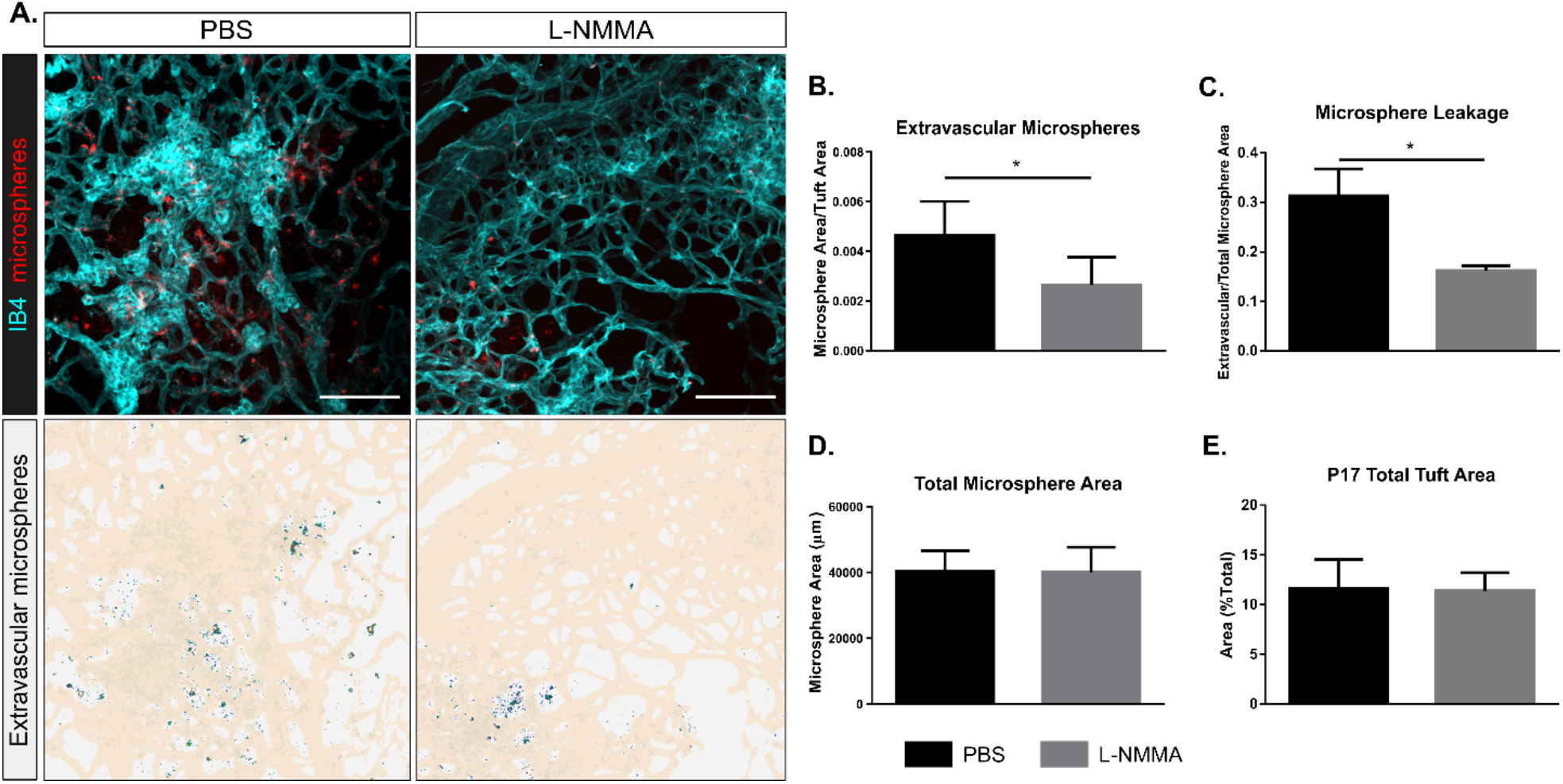
Decreased leakage from retinal vascular tufts after single dose L-NMMA treatment. A. Representative images of tufts from *Nos3*^*+/+*^ (wild type) mice treated with PBS or L-NMMA (60 μg/g body weight) 24 hrs before tail-vein injection of 25 nm microspheres. Retinas were immunostained for isolectin B4 (IB4; cyan), microspheres (red) appear in and around the tufts. Scale bar = 100 μM. Lower panels show leakage maps. Heat mapped dots that do not overlap with vessels (beige) are considered extravascular. B. Quantification of the average area of extravascular microspheres normalized to IB4 area. C. Quantification of the average area of extravascular microspheres normalized to total microsphere area. D. Quantification of average total microsphere area in PBS or L-NMMA-treated wildtype mouse retinas. E. Quantification of tuft area normalized to total vascular area. Mean ±S.E.M. n = 5 (PBS) and 5 (L-NMMA) treated mice. * = p < 0.05; t-test. Figure 5 – source data 1. Raw data on retina vascular parameters and body weights from *Nos3*^*+/+*^ mice treated with L-NMMA and injected with microspheres

## Discussion

The results presented here show that eNOS/NO can modulate endothelial barrier function to exacerbate vascular hyperpermeability in retinopathy by a direct effect on endothelial junctions (Figure 6). Historically, eNOS-generated NO is implicated in the regulation of vascular permeability by inducing the relaxation of perivascular smooth muscle cells. NO produced in endothelial cells diffuses across the vascular wall and activates soluble guanylate cyclase leading to protein kinase G activation in smooth muscle cells, lowering cellular Ca^2+^ and promoting vascular relaxation, increased blood flow and reduced blood pressure (Surks et al., 1999). Indeed, vessel dilation is a part of the tissue deterioration seen in diabetic retinopathy (Bek, 2013; Grimm & Willmann, 2012). However, there are also indications for a direct role for eNOS and NO in endothelial cells, as constitutive eNOS deficiency inhibits inflammatory hyperpermeability in mouse cremaster muscle treated with platelet-activating factor (Hatakeyama et al., 2006). Moreover, NO can regulate phosphorylation of VE-cadherin in adherens junctions *in vitro* in microvascular endothelial cell cultures (Di Lorenzo et al., 2013). It is likely that these multifaceted effects of eNOS/NO are differently established in different vessel types. eNOS/NO-dependent vessel dilation is dependent on the vascular smooth muscle cell coverage in arterioles and arteries, in contrast, adherens junction stability affects mainly the postcapillary venular bed (Orsenigo et al., 2012). In the skin, prevenular capillaries and postcapillary venules, with sparse vSMC coverage, respond to VEGFA with increased paracellular permeability while arterioles/arteries do not (Honkura et al., 2018). These distinctions are important as exaggerated and chronic VEGFA-driven paracellular permeability in disease leads to edema and eventually to tissue destruction (Nagy, Dvorak, & Dvorak, 2012). Therefore, we explored the consequences of NO-deficiency using a mutant eNOS mouse model, with a serine to alanine exchange at 1176, as well as treatment with the NO-inhibitor L-NMMA, to examine how attenuating NO-production affects VEGFA-dependent pathological angiogenesis in ocular disease.

**Figure 6.**
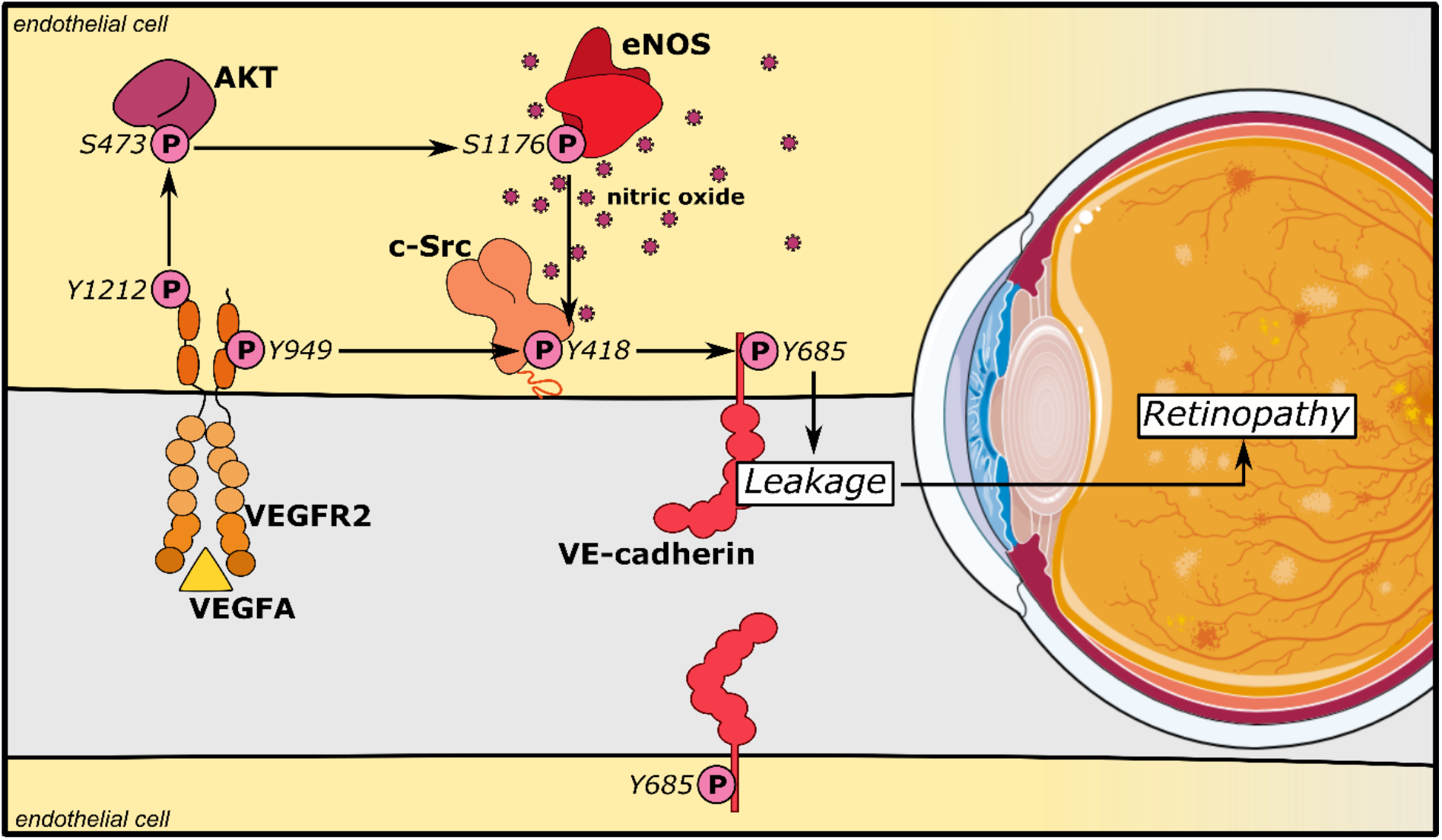
eNOS/NO modulates VE-cadherin Y685 phosphorylation via c-Src in a VEGFA/VEGFR2 dependent manner. VEGFA through VEGFR2 and its phosphosite Y1212 induces a chain of consecutive reactions in endothelial cells: phosphorylation of AKT at S473 and eNOS at S1176. The VEGFR2 phosphosite Y949 mediates phosphorylation of c-Src at Y418 and of VE-cadherin at Y685. Combined, these activating phosphorylation reactions disrupt the vascular barrier by dissociating VE-cadherin’s homophilic interactions, resulting in macromolecular leakage. eNOS/NO exacerbates this damage via an interaction with c-Src to enhance VE-cadherin Y685 phosphorylation and internalization.

Our data shows that while the attenuation of eNOS S1176 phosphorylation was dispensable for vascular development in the retina and for endothelial survival during hyperoxia, growth of pathological vessel tufts in the subsequent phase of relative hypoxia was suppressed. Similarly, treatment with L-NMMA during the hypoxic phase reduced growth of vascular tufts in the retina. In agreement with earlier literature, we conclude that eNOS activity and NO formation influences pathological angiogenesis in the eye (Ando et al., 2002; Brooks et al., 2001; Edgar, Gardiner, van Haperen, de Crom, & McDonald, 2012). Of note, tufts that were established in the *Nos3*^*S1176A/S1176A*^ mice leaked less, in spite of similar levels of VEGFA being produced as in the wildtype retina. In an attempt to mimic the clinical situation, we treated P16 mice with established vascular tufts with a single dose of L-NMMA. At the examination 24 h later, tuft area remained unaffected while the leakage of 25 nm microspheres was reduced by 50-60%. Whether leakage regulation is separable from the regulation of growth of new vessels has been a matter of debate. Pathological angiogenesis in the retina is intimately associated with leakage and edema (Smith et al., 2020). Exactly how junction stability plays a role in the neoangiogenic process is unclear. However, leakage and the production of a provisional matrix is postulated to be a prerequisite for the growth of angiogenic sprouts (reviewed in (Nagy et al., 2012)). With the effects of the L-NMMA intervention treatment shown here, we can conclude that these responses indeed can be separated, at least in the short term.

L-NMMA, and its analog L-NAME, induce vasoconstriction by halting the NO/soluble guanylyl cyclase signaling pathway and consequent vasodilation (reviewed in (Ahmad et al., 2018; Thoonen, Sips, Bloch, & Buys, 2013)). Thus, a daily intake of L-NAME in rats (40-75 μg/g/day for 4 – 8 weeks) leads to vasoconstriction (Ribeiro, Antunes, de Nucci, Lovisolo, & Zatz, 1992; Simko et al., 2018; Vrankova, Zemancikova, Torok, & Pechanova, 2019; Zanfolin et al., 2006). NOS inhibitors have also been used clinically, for example by administration of L-NMMA to increase the mean arterial pressure in cardiogenic shock. In a typical regimen, 1mg.kg^−1^.hr^−1^ of L-NMMA is administered over 5 hours (Cotter et al., 2000). While we cannot exclude an effect on vasoconstriction with the single IP injection of L-NMMA used here (60 μg/g), blood flow appeared unaffected as equal amounts of microspheres arrived in the retinal vasculature, while leakage into the extravascular space was substantially reduced in the L-NMMA treated mice compared to the controls.

Mechanistically, our results place c-Src downstream of eNOS activity. c-Src has also been placed upstream of eNOS by c-Src’s regulation of the PI3K-AKT pathway, of importance for eNOS activation (Haynes et al., 2003). VEGFR2-induced activation of c-Src has however been mapped to the pY949 residue in VEGFR2 (Li et al., 2016) while activation of Akt is dependent on Y1212 (Testini et al., 2019). In VEGFA-stimulated endothelial cells, c-Src is implicated in phosphorylation of VE-cadherin at Y685 and potentially other tyrosine residues, to induce vascular permeability (Orsenigo et al., 2012; Wallez et al., 2007). The fact that the mutant *VEC-Y685F* mice in which the Y685 residue has been replaced with a non-phosphorylatable phenylalanine, were resistant to L-NMMA inhibition, indicates that the VEGFR2 Y1212/eNOS/NO pathway acts on adherens junction through c-Src (Li et al., 2016). Combined, our data support a model in which eNOS phosphorylation at S1176 is required for the activation of c-Src, which in turn phosphorylates VE-cadherin at Y685, inducing transient disintegration of adherens junctions and increased paracellular permeability (Figure 6). These *in vivo* results provide meaningful mechanistic and therapeutic insight into retinal diseases accompanied by excessive permeability, such as diabetic retinopathy, age-related macular degeneration and retinopathy of prematurity (Antonetti, Klein, & Gardner, 2012; Cunha-Vaz, Bernardes, & Lobo, 2011).

A considerable challenge in analysis of c-Src activity is the close structural relatedness between the kinase domains of c-Src and the Src family members (SFKs) Yes and Fyn (Sato et al., 2009), which are also expressed in endothelial cells. The amino acid sequence covering the activating tyrosine is entirely conserved between these three cytoplasmic tyrosine kinases such that the antibodies against pY418 c-Src in fact reacts with all three SFKs. Thus, pY418 immunostaining of wildtype and *Nos3*^*S1176A/S1176A*^ retinas after OIR failed to reveal a dependence on eNOS catalytic activity for activation of c-Src, in agreement with the findings of Di Lorenzo et al. (Di Lorenzo et al., 2013). However, combining oligonucleotide-linked secondary antibodies reacting with c-Src protein and pY418 Src in a PLA on isolated endothelial cells demonstrated a critical role for eNOS activity on accumulation of active, pY418-positive c-Src at endothelial junctions. The question remains how eNOS/NO influence c-Src activity? NO can couple to cysteine thiols to form S-nitroso-thiols, which may affect the folding and function of the target protein. In accordance, Rahman et al. demonstrated S-nitrosylation of c-Src at the kinase domain cysteine 498, correlating with increased c-Src activity (Rahman et al., 2010).

Patients with diabetic retinopathy have elevated levels of NO in the aqueous humor and particular eNOS polymorphisms are associated with protection or increased risk for diabetic retinopathy (for a review, see (Opatrilova et al., 2018)) and macular edema (Awata et al., 2004). Our results show that pharmacological inhibition of NO production also in established disease can prevent vascular leakage. Thus, NO inhibitors applied in combination with anti-VEGF therapy could possibly be delivered in low but still efficient doses. Local delivery of NOS inhibitors may be needed to avoid drawbacks with systemic delivery such as hypertension, or any other vasoconstriction-associated adverse events.

In conclusion, the data presented here establish a critical role for eNOS in endothelial cells, regulating c-Src activation downstream of VEGFA/VEGFR2, and thereby, in VE-cadherin-regulated endothelial junction stability and vascular leakage in retinal pathology.

## Materials and methods

### Key Resources Table

| Reagent type | Designation | Source/reference | Identifiers | Information |
| --- | --- | --- | --- | --- |
| *Strain; strain background (Mus musculus)* | Nos3<sup>+/+</sup> | 10.1016/j.bbrc.2012.12.110 |  | C57BL/6J |
| *Strain; strain background (Mus musculus)* | Nos3<sup>S1176A/S1176A</sup> | 10.1016/j.bbrc.2012.12.110 |  | C57BL/6J |
| *Strain; strain background (Mus musculus)* | VEC-WT | DOI: 10.1038/ni.2824 |  | C57BL/6J |
| *Strain; strain background (Mus musculus)* | VEC-Y685F | DOI: 10.1038/ni.2824 |  | C57BL/6J |
| *Antibody* | Rat anti-VE-cadherin | BD Biosciences | Catalogue no. 555289<br>RRID:AB_395707 | Dilution; 1:100 |
| *Antibody* | Rabbit anti-VE-cadherin pY685 | DOI: 10.1038/ncomms2199 |  | Dilution; 1:50 |
| *Antibody* | Goat anti-VE-cadherin | R&D systems | Catalogue no. AF1002<br>RRID:AB_2077789 | Dilution; 1:200 |
| *Antibody* | Mouse anti-eNOS | Abcam | Catalogue no. ab76198<br>RRID:AB_1310183 | Dilution; 1:1000 |
| *Antibody* | Mouse anti-Src GD11 | Merck Millipore | Catalogue no. 05-184<br>RRID:AB_2302631 | Dilution; 1:200 |
| *Antibody* | Rabbit anti-c-Src pY418 | Invitrogen | Catalogue no. 44-660G<br>RRID:AB_1500523 | Dilution; 1:100 |
| *Antibody* | Mouse anti-α-tubulin | Sigma-Aldrich | Catalogue no. T9026 | Dilution; 1:1000 |
|  |  |  | RRID:AB_477593 |  |
| *Antibody* | Mouse anti-eNOS pS1177 | BD Biosciences | Catalogue no. 612392<br>RRID:AB_399750 | Dilution; 1:1000 |
| *Antibody* | Rat anti-CD31 | BD Biosciences | Catalogue no. 553370<br>RRID:AB_394816 | Dilution; 1:200 |
| *Antibody* | Rabbit anti-ERG | Abcam | Catalogue no. Ab92513<br>RRID:AB_2630401 | Dilution; 1:200 |
| *Antibody* | Donkey anti-Rabbit IgG | Thermo Fisher Scientific | Catalogue no. A-31572<br>RRID:AB_162543 | Dilution; 1:500 |
| *Antibody* | Donkey anti-Rat IgG | Thermo Fisher Scientific | Catalogue no. A-21208<br>RRID:AB_141709 | Dilution; 1:500 |
| *Antibody* | Donkey anti-Goat IgG | Thermo Fisher Scientific | Catalogue no. A-11055 | Dilution; 1:500 |
| *Antibody* | Donkey anti-Mouse IgG, HRP | Thermo Fisher Scientific | Catalogue no. A-16011 | Dilution; 1:1000 |
| *Kit* | Griess Assay (Nitrate/Nitrite colorimetric assay kit) | Cayman Chemical | Catalogue no. 780001 |  |
| *Reagent* | CD31 microbeads, mouse | Miltenyi Biotec | Catalogue no. 130-097-418 |  |
| *Reagent* | Nω-Methyl-L-arginine acetate salt (L-NMMA) | Sigma-Aldrich | Catalogue no. M7033 | |
| *Reagent* | Amersham ECL Prime Western Blotting Detection | GE healthcare | Catalogue no. RPN2232 |  |
| *Other* | Nos3 primer | Thermo Fisher Scientific | Mm00435217_m1 |  |
| Other | Nos2 primer | Thermo Fisher Scientific | Mm00440502_m1 |  |
| Other | Vegfa primer | Thermo Fisher Scientific | Mm00437306_m1 |  |
| Other | TBP primer | Thermo Fisher Scientific | Mm01277042_m1 |  |
| Other | UBC primer | Thermo Fisher Scientific | Mm02525934_g1 |  |
| Other | VEGFA<sub>164</sub>, recombinant, mouse | R&D systems | Catalogue no. 493-MV/CF |  |
| Other | Fluoro-Max Dyed Green Aqueous Fluorescent Particles | Thermo Fisher Scientific | Catalogue no. G25 |  |
| Other | Duolink In Situ PLA Probe anti-Rabbit MINUS | Sigma-Aldrich | Catalogue no. DUO92005 |  |
| Other | Duolink In Situ PLA Probe anti-Mouse PLUS | Sigma-Aldrich | Catalogue no. DUO92001 |  |
| Other | Duolink In Situ Detection Reagent (Orange) | Sigma-Aldrich | Catalogue no. DUO92007 |  |
| Other | Hoechst 33342 | Thermo Fisher Scientific | Catalogue no. H3570 | Dilution; 1:1000 |
| Other | Alexa Fluor 488-Isolectin B4 | Sigma-Aldrich | Catalogue no. I21411<br>RRID:AB_2314662 | Dilution; 1:200 |
| Other | Alexa Fluor 647-Isolectin B4 | Sigma-Aldrich | Catalogue no. I32450<br>RRID:SCR_014365 | Dilution; 1:200 |
| Software | ImageJ | NIH, Bethesda, MD, USA | RRID:SCR_003070 |  |
| Software | GraphPad Prism | GraphPad | RRID:SCR_002798 |  |

### Animal studies

*Nos3*^*S1176A/S1176A*^ mice on a C57Bl/6J background have been described (Schleicher et al., 2009). *VEC*^*Y685FY685F*^ mice, also on C57Bl/6J background were generated using site-directed mutagenesis on a wildtype human VE-cadherin cDNA construct to create the Y685 to F685 mutation (Wessel et al., 2014). Both strains were maintained by crossing heterozygous mice. Wildtype C57BL/6J mice (Jackson Laboratory) and the Y685F strain were treated, when indicated, with Nω-Methyl-L-arginine acetate salt (L-NMMA; Sigma-Aldrich) in PBS, 20 μg/g body weight, by intraperitoneal injection from postnatal (P) day 12 to P16.

Wildtype mice were also treated, when indicated, with a single dose of L-NMMA in PBS, 60 μg/g body weight, by intraperitoneal injection on P16. These mice were then used to perform microsphere assays on P17.

Mouse husbandry and OIR challenge took place at Uppsala University, and the Local Ethics committee approved all animal work for these studies Animal handling was in accordance to the ARVO statement for the Use of Animals in Ophthalmologic and Vision Research. All animal experiments were repeated in individually at least three times (biological repeats).

### Oxygen-induced retinopathy (OIR)

A standard oxygen-induced retinopathy model was used (Connor et al., 2009). Briefly, each litter of pups was placed, along with the mother, into a chamber maintaining 75.0% oxygen (ProOx 110 sensor and AChamber, Biospherix, Parish, NY) from P7-P12, when they returned to normal atmosphere (~21% oxygen) until P17 (termination). The lactating mother was removed each day, P8-P11, for 2 hours, to prevent oxygen toxicity. At P17, pups were weighed and sacrificed. Eyes were enucleated and fixed in 4% paraformaldehyde (PFA) at room temperature for 30 minutes. See source data files for data on neoangiogenic tufts, avascular area and body weights at P17. No mice were excluded from analysis.

### Quantification of avascular area and neovascular tufts

Avascular area and neovascular tuft formation were determined by immunostaining retinas followed by imaging (Leica SP8 confocal microscope) and analysis. Quantification of total vascularized area, central avascular area, and tuft area was performed by outlining images manually in ImageJ (NIH, Bethesda, MD). Using a tilescan of IB4 staining for the entire retina, the freehand selection tool was used to demarcate the vascular front, creating an ROI (region of interest) for the total vascularized area. The freehand selection tool was used to outline IB4 positive vessels from neovascular tufts (regions with disorganized dilated vessels). The ROIs for tufts were merged into a single ROI corresponding to the total neovascular tuft area for each retina. The tuft area normalized to the total vascularized area was reported as a percentage of the total retina that contained tufts. Similarly, the avascular region was determined using the freehand selection tool to outline the central avascular regions. Regions where the superficial layer of capillaries was absent were determined and merged to form a single ROI corresponding to the entire avascular region for each retina. The avascular area normalized to the total vascularized area was reported as a percentage of the total retina that was still avascular. The researcher was blinded to the genotype of the sample when performing quantifications

### Microsphere assay

For microsphere extravasation experiments, mice at P17 were briefly warmed under heat lamp to dilate tail veins before injection of microspheres (1% solution of 25 nm fluorescent microspheres; 50 μl per mouse into the tail vein; ThermoFisher). Microspheres circulated for 15 minutes. To remove blood and microspheres from the retinal vessels, mice were perfused with room temperature phosphate-buffered saline (PBS) containing calcium chloride and magnesium chloride using a peristaltic pump for 2 min, under full isoflurane-anesthesia. The eyes were then enucleated and fixed in 4% paraformaldehyde (PFA) at room temperature for 30 minutes before dissection and mounting for microscopy (Leicas SP8 confocal microscope, 63x objective).

Using ImageJ software, the microsphere channel and IB4-vessel channel (488 for Green fluorescence and 647 for IB4) were adjusted with threshold (Huang for IB4 and Triangle for FITC) for each channel. Extravasated microsphere area was calculated by measuring the signal in the green fluorescence channel after removing any signal contained within the ROI corresponding to the IB4-positive area. The Analyze Particles function was employed to quantify the microspheres. A lower limit of 10 pixels was selected to distinguish the microsphere signal from background noise. The mean area density for each group of mice was calculated from the median value of all images of the eyes of each mouse (Fuxe et al., 2011). To quantify leakage based on microscopic images, the amount of tracer extravasation was normalized to blood vessel density. The researcher was blinded to the genotype of the sample when performing quantifications. See source files for data on neoangiogenic tufts and body weights at P17. No mice were excluded from analysis.

### Endothelial cell isolation from mouse lung

Mouse lungs were harvested from pups (age; P8 - P10), minced and digested in 2 mg/ml collagenase type I (385 U/mg; Worthington) in PBS with Ca^2+^/Mg^2+^ for 1 h at 37°C. Cells were then isolated using CD31 Microbeads and MACS cell isolation equipment and reagents (Miltenyi Biotec). The cells were seeded at 3×10^5^ cells/mL and cultured in MV2 medium with supplements and serum (PromoCell).

### Griess Reagent Assay

Isolated endothelial cells were seeded at 3×10^5^ cells/mL into 6 cm cell culture plates and allowed to adhere at 37°C and 5% CO_2_ overnight. After 24 hrs a Griess Assay was performed (nitrate/nitrite colorimetric assay, Cayman chemical) according to the manufacturer’s instruction. Once complete, the cells were lysed in 1% [w/v] NP 40, 25 mM Tris HCl pH 7.6, 0.1% SDS, 0.15 M NaCl, 1% sodium deoxycholate, 1x Protease Inhibitor Cocktail (Roche) and concentration of nitrate/nitrite was normalised against cell protein concentration, measured using the BCA protein detection kit (ThermoFisher).

### Proximity Ligation Assay (PLA)

Isolated endothelial cells, serum starved at 37°C in MV2 medium (containing no growth factors) 3 hrs before stimulation with VEGFA_164_ (50 ng/ml; R&D Systems), followed by fixation in 3% PFA for 3 min, permeabilized in 0.1% Triton X-100 for 3 min, and postfixed in 3% PFA for 15 min. Samples were blocked in Duolink blocking buffer for 2 hours at 37°C and used for PLA. The Duolink protocol (Sigma-Aldrich) was followed using anti-phospho-Src Tyr 418 (Invitrogen) and total c-Src (Merck Millipore) antibodies, and oligonucleotide-linked secondary antibodies, denoted PLUS and MINUS probes, followed by the detection of reactions with fluorescent probes. Upon completion of the PLA protocol, cells were counterstained with antibodies against VE-cadherin (R&D systems), and Hoechst 33342 (ThermoFisher) to detect nuclei. Only cells positive for VE-cadherin were imaged and analyzed. To determine c-Src p418 association with VE-cadherin, a mask of the VE-cadherin channel was created and only points that aligned completely within the VE-cadherin mask were counted and expressed against the area of VE-cadherin per field of view (ImageJ software, NIH). As a technical control for each experiment, the same procedure was performed with the omission of either of the primary antibodies, or the PLUS/MINUS probes.

### Immunofluorescent staining

Whole mount immunostaining was performed on PFA-fixed retinas incubated in blocking buffer for 2 hours (Buffer b; bovine serum albumin (BSA)/2% fetal calf serum (FCS)/0.05% Na-deoxycholate/0.5% Triton X-100/0.02% Na Azide in PBS). Incubation with primary antibodies over night at 4°C on a rocking platform was followed by incubation with secondary antibodies overnight at 4°C. Retinas were mounted on slides using Fluormount G. Images were taken by Leica SP8 confocal microscope and acquired with the 10x or 63x objective. Processing and quantification of images was done with ImageJ software (NIH). Quantification in the retina of total vascularized area, central avascular area, and area covered by neovascular tufts was performed by outlining images manually in ImageJ. Avascular area and tuft area were normalized to the total vascularized area of the retina.

### Quantitative real-time PCR

RNA from retinas were purified using RNeasy Kit (Qiagen). One microgram of RNA was reverse transcribed using SuperScript III (Invitrogen) and quantitative PCR were assayed using Mus musculus primers against *Vegfa* (Mm00437306_m1, ThermoFisher), *Nos3* (Mm00435217_m1) and *Nos2* (Mm00440502_m1). The expression levels were normalized against TATA binding protein (*TBP)* Mus musculus (Mm01277042_m1,ThermoFisher) and Ubiqutin C (*UBC*) Mus musculus (Mm02525934_g1, ThermoFisher).

### Cells culture and treatment

Human retinal microvascular endothelial cells (HRMECs; Cell Systems, #ACBRI 181) were cultured in a complete classic medium kit with serum and CultureBoost (Cell Systems, #4Z0–500). The cells were used and passaged in 10 cm cell culture plates coated with attachment factor, between passages 5–10 for all experiments. All cells were serum starved for 3 hrs at 37°C in MV2 medium (containing no growth factors) before stimulation. Recombinant mouse VEGFA_164_ (R&D Systems), was used at 50 ng/ml for in vitro analyses. L-NMMA (1 mM in PBS) was administrated 1 hr before stimulation with VEGFA.

### Western blot

Cells were lysed in 1% [w/v] NP 40, 25 mM Tris HCl pH 7.6, 0.1% SDS, 0.15 M NaCl, 1% sodium deoxycholate, 1x Protease Inhibitor Cocktail (Roche), 1 mM Na_3_VO_4_ (Sigma), and centrifuged at 21,100 g for 10 min. Protein concentration was measured with the BCA protein detection kit (ThermoFisher). Proteins were separated on a 4-12% BisTris polyacrylamide gel (Novex by Life Technologies) and transferred to an Immobilon-P PVDF membrane (Millipore) using the Criterion Blotter system (BioRad). The membrane was blocked with 3-5% skim milk in Tris-buffered saline (TBS; 0.1% Tween). For phosphotyrosine antibodies blocking was done in 5% BSA in TBS, 0.1% Tween. The membrane was incubated with first antibodies overnight at 4°C. Membranes were then washed in TBS, 0.1% Tween and incubated with horseradish peroxidase (HRP)-conjugated secondary anti-mouse antibody (1:10,000; Invitrogen) in 3-5% skim milk, respectively, followed by final wash in TBS, 0.1% Tween and development using ECL prime (GE Healthcare). Luminescence signal was detected by the ChemiDoc MP system (BioRad) and densitometry performed using Image Lab software (ver 4, BioRad)

### Antibodies

Retinal vasculature was immunostained with directly-conjugated Alexa Fluor 488-Isolectin B4 (1:200; Sigma, I21411) or Alexa Fluor 647-Isolectin B4 (1:200; Sigma, I32450). EC junctions and phosphorylated VE-cadherin were detected with Anti-VE-cadherin antibody (1:100; BD, Rat, 555289) and affinity purified rabbit antibodies against VE-cadherin pY685; a kind gift from Prof. Elisabetta Dejana, Uppsala University/IFOM Milano^18^. For proximity ligation assays VE-cadherin was detected using mouse VE-cadherin antibody (1:200, R&D systems, Goat, AF1002). c-Src was detected using anti-Src (GD11 clone) antibody (1:200, Merck Millipore, Mouse, 05-184). Phosphorylated c-Src was assessed using anti-phospho-Src Tyr 418 antibody (1:100, Invitrogen, Rabbit, 44-660G). Nuclei were detected using Hoechst 33342 (1:1000, ThermoFisher, H3570). For immunoblotting, the following antibodies were used as primaries: mouse-anti-αtubulin (1:1000, Sigma, T9026), mouse anti-eNOS (1:1000, Abcam, ab76198), mouse anti-eNOS pS1177 (1:1000, BD, 612392). Secondary antibody: Amersham ECL Mouse IgG, HRP-linked whole Ab (from sheep) (1:10,000, GE Healthcare, NA931). Detection: Amersham ECL prime Western blotting detection reagent (GE healthcare, RPN2232).

### DAF FM DA assay

Intracellular NO was measured in real time using the NO-specific fluorescence probe DAF-FM DA solution (Sigma Aldrich). DAF-FM DA diffuses freely across the membrane, and is hydrolyzed by intracellular esterases, resulting in the formation of DAF-FM. Intracellular DAF-FM reacts with the NO oxidation product N_2_O_2_, which generates the stable highly fluorescent derivative DAF-FM triazole. Cells were washed with modified HEPES Buffer (20 mM HEPES buffer (Gibco) with 5 mM glucose, 50 μM L-Arginine and 0.1% BSA, pH7.0-7.4), incubated with 5 μM DAF-FM DA in modified HEPES buffer for 30 min at room temperature, washed again and finally incubated in modified HEPES buffer for 30 min at 37°C in the absence or presence of 1 mM L-NMMA. Fluorescence (emission wavelength, 485 nm; excitation wavelength, 538 nm) was measured at 37°C from 1 to 10 min using a fluorescence microtiter plate reader (Synergy HTX Multi-Mode Reader, BioTek, USA). eNOS activity was expressed as the VEGFA-dependent increase in fluorescence per μg of cellular protein. To determine the cellular protein content the same cells were lysed in 1% (v/v) Triton X-100 and analyzed for protein content with the BCA protein detection kit. DAF-FM DA experiments were repeated three times. Within each experiment, four wells were used for each NO measurement.

### Statistical analysis

Statistical analysis was performed using GraphPad Prism 6 (GraphPad). An unpaired Student’s T test was used to compare means among two experimental groups. Two-way ANOVAs were performed when two factors were involved, for example, treatment and genotype. Multiple comparisons post hoc tests were chosen based on how many group comparisons were made. In all analyses p < 0.05 was considered a statistically significant result. Values shown are the mean, with standard error of the mean (S.E.M.) used as the dispersion measure. Biological replicates refer to individual mice/samples in a single experiment. Separate/individual experiments refer to experiments done at different times/days with independently generated material. A statistical method of sample size calculation was not used during the study design.

## Abbreviations

eNOS: endothelial nitric oxide synthase
L-NMMA: Nω-Methyl-L-arginine acetate
Nos3: gene designation for murine endothelial nitric oxide synthase 3
OIR: oxygen-induced retinopathy
PLA: proximity ligation assay
VEGFA: vascular endothelial growth factor A
VE-cadherin: vascular endothelial-cadherin

## Acknowledgements

Acknowledgement: We gratefully acknowledge the expert assistance of Pernilla Martinsson, Uppsala University.

Sources of funding: This study was supported by the Swedish Cancer foundation (CAN2016/578), the Swedish Research Council (2015-02375), the Knut and Alice Wallenberg foundation (KAW 2015.0030) and a Fondation Leducq Transatlantic Network of Excellence Grant in Neurovascular Disease (17 CVD 03). KAW also supported LCW with a Wallenberg Scholar grant (2015.0275). WCS was supported by Grants R35 HL139945, P01 HL1070205, AHA MERIT Award.

## Competing Interests

Disclosure: The authors declare no competing interests.

---

**Figure 1 – supplement figure 1.**
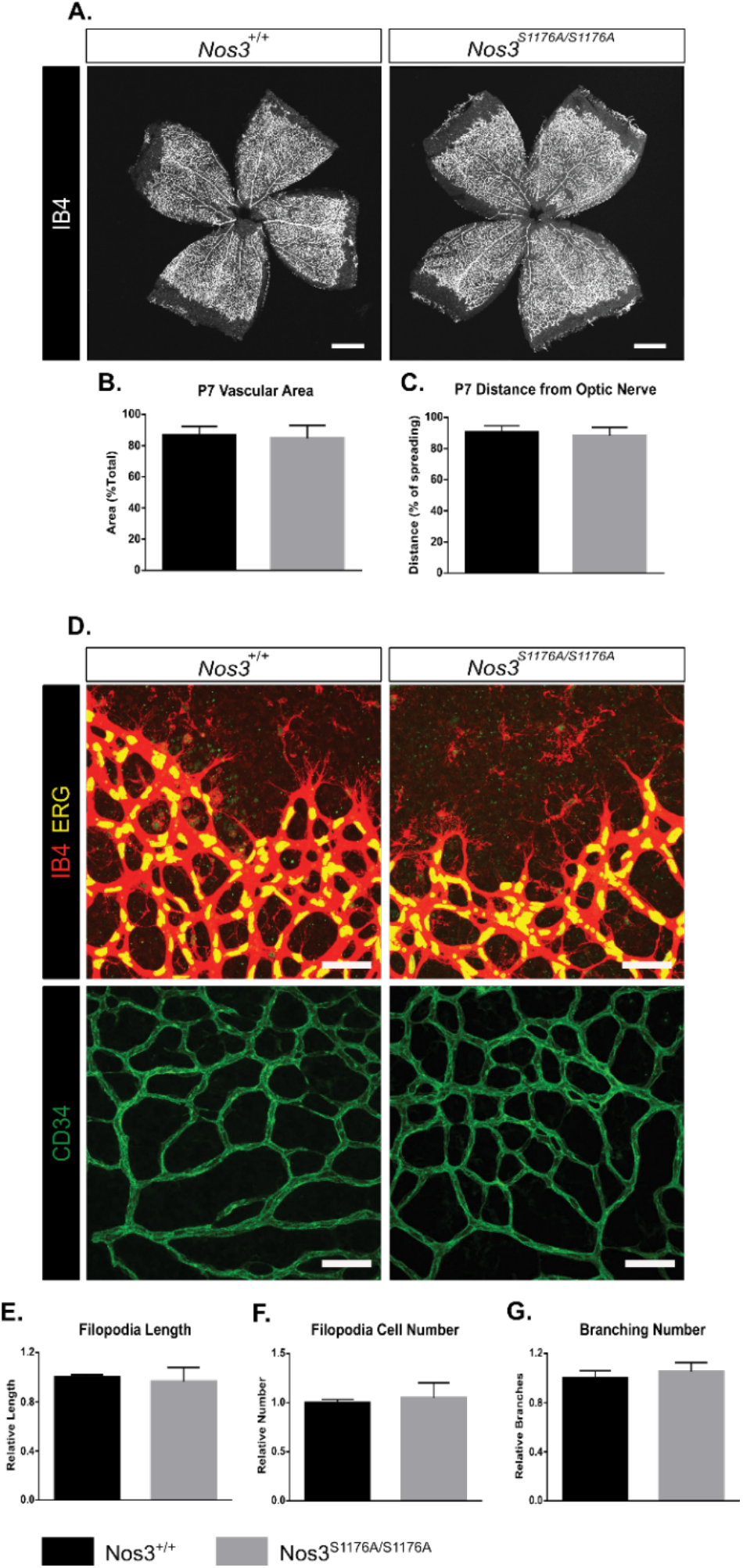
Postnatal development of *Nos3*^*+/+*^ and *Nos3*^*S1176A/S1176A*^ retinal vasculature. A. Representative images of Nos3^+/+^ and Nos3^S1176A/S1176A^ retinas collected at P7, stained with isolectin B4 (IB4). B, C. Quantification of vascular area (B) and outgrowth from the optic nerve (C) at P7 in *Nos3+/+* and *Nos3S1176A/S1176A* pups. D. Representative images of the vessel front from whole mount retinas collected at P7 stained with Isolectin-B4 (IB4, red in upper and green in lower panels) and ERG (yellow) to visualise vessel outgrowth and tip cells in *Nos3*^*+/+*^ and *Nos3*^*S1176A/S1176A*^ retinal vasculature. Scale bar = 50 μm. E-G. Filopodia length (E), tip cell number (F), branching points (G) in *Nos3*^*+/+*^ and *Nos3*^*S1176A/S1176A*^ retinas at P7. Mean ± S.E.M. n=3-4 mice, 5-6 images/mouse; t-test.

**Figure 1 – supplement figure 2.**
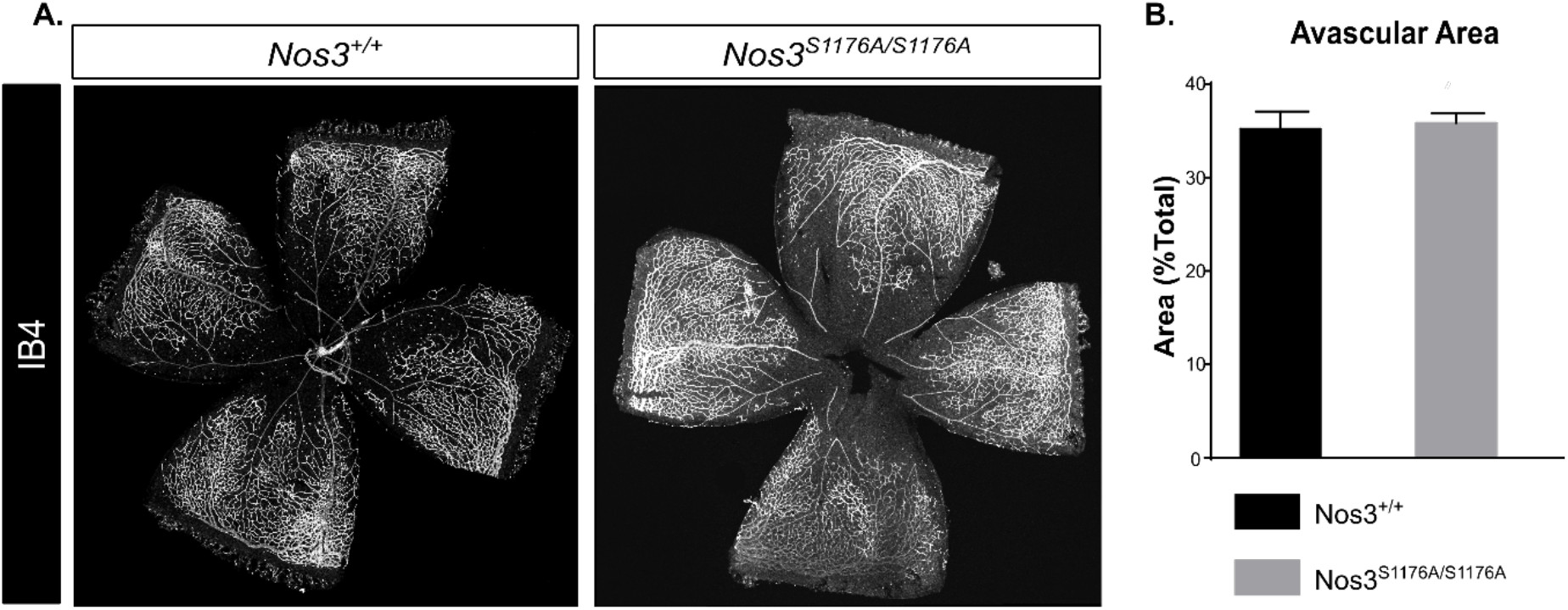
Retina development in *Nos3*^*+/+*^ and *Nos3*^*S1176A/S1176A*^ P12 pups. A. Representative images of whole mount *Nos3*^*+/+*^ and *Nos3*^*S1176A/S1176A*^ retinas collected at P12 after the vessel destruction phase of OIR and before vessel regrowth, stained with isolectin B4 (IB4). Scale bar = 500 μm. B. Avascular area in *Nos3*^*+/+*^ and *Nos3*^*S1176A/S1176A*^ retinas at P12 after OIR. n = 4 (*Nos3*^*+/+*^) and 5 (*Nos3*^*S1176A/S1176A*^) mice, 3 independent experiments; t-test.

**Figure 1 – supplement figure 3.**
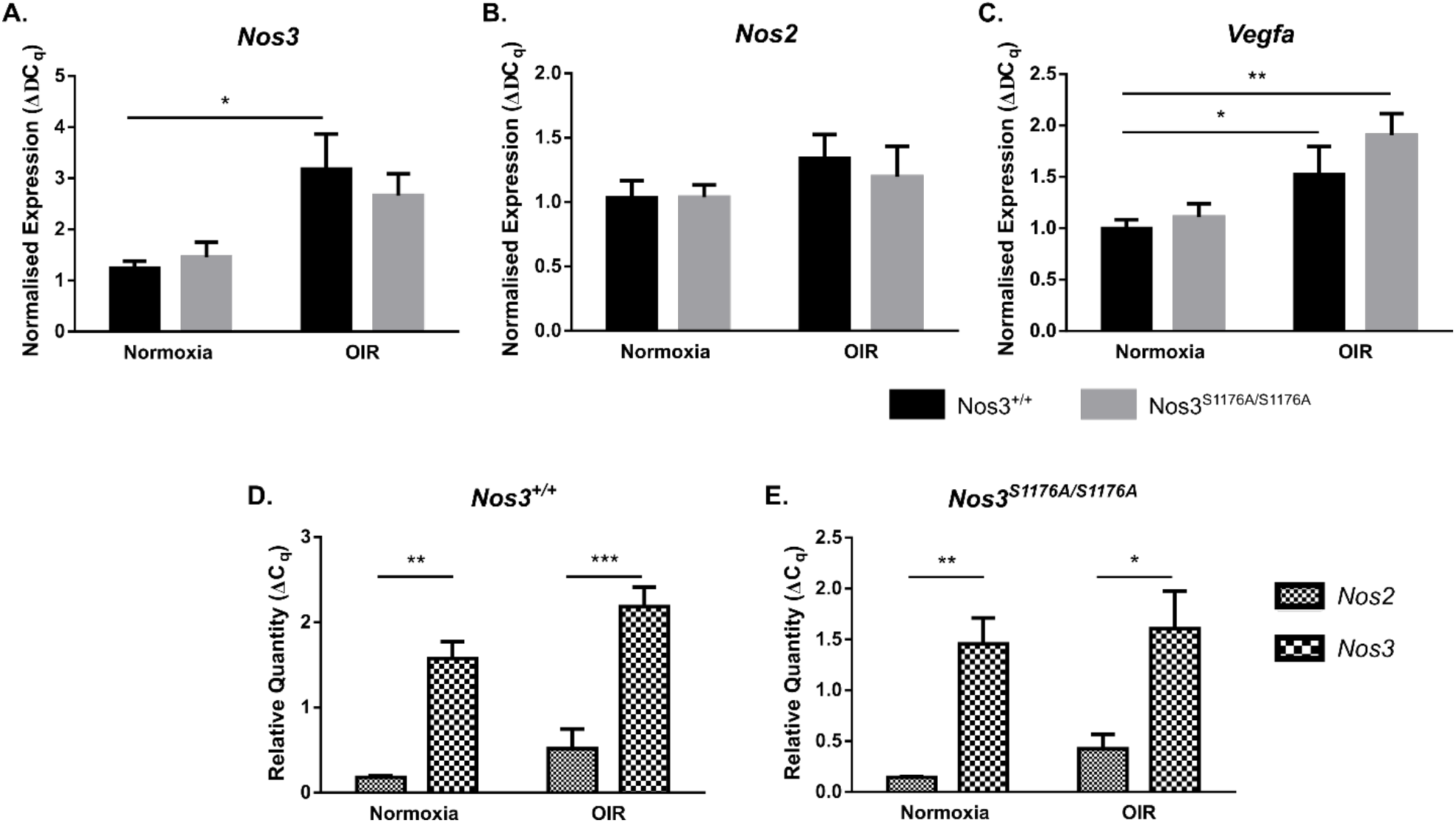
Expression of Nos2, Nos3 and Vegfa in *Nos3*^*+/+*^ and *Nos3S1176A/S1176A* retinas. A-C. qPCR of *Nos3 (A)*, *Nos2 (B)* and *Vegfa* (C) expression in P17 normoxic and OIR-challenged *Nos3*^*+/+*^ and *Nos3*^*S1176A/S1176A*^ mouse retinas. D,E. Relative quantities of *Nos2* and *Nos3* compared against standard curves of *TBP* and *UBC* in *Nos3*^*+/+*^ and *Nos3*^*S1176A/S1176A*^ retinas. Mean ±S.E.M. n = 5 (*Nos3*^*+/+*^) and 5 (*Nos3*^*S1176A/S1176A*^) mice. *, **, *** = p < 0.05, 0.01, 0.001; two-way ANOVA, Sidak’s multiple comparison test.

**Figure 2 – supplement figure 1.**
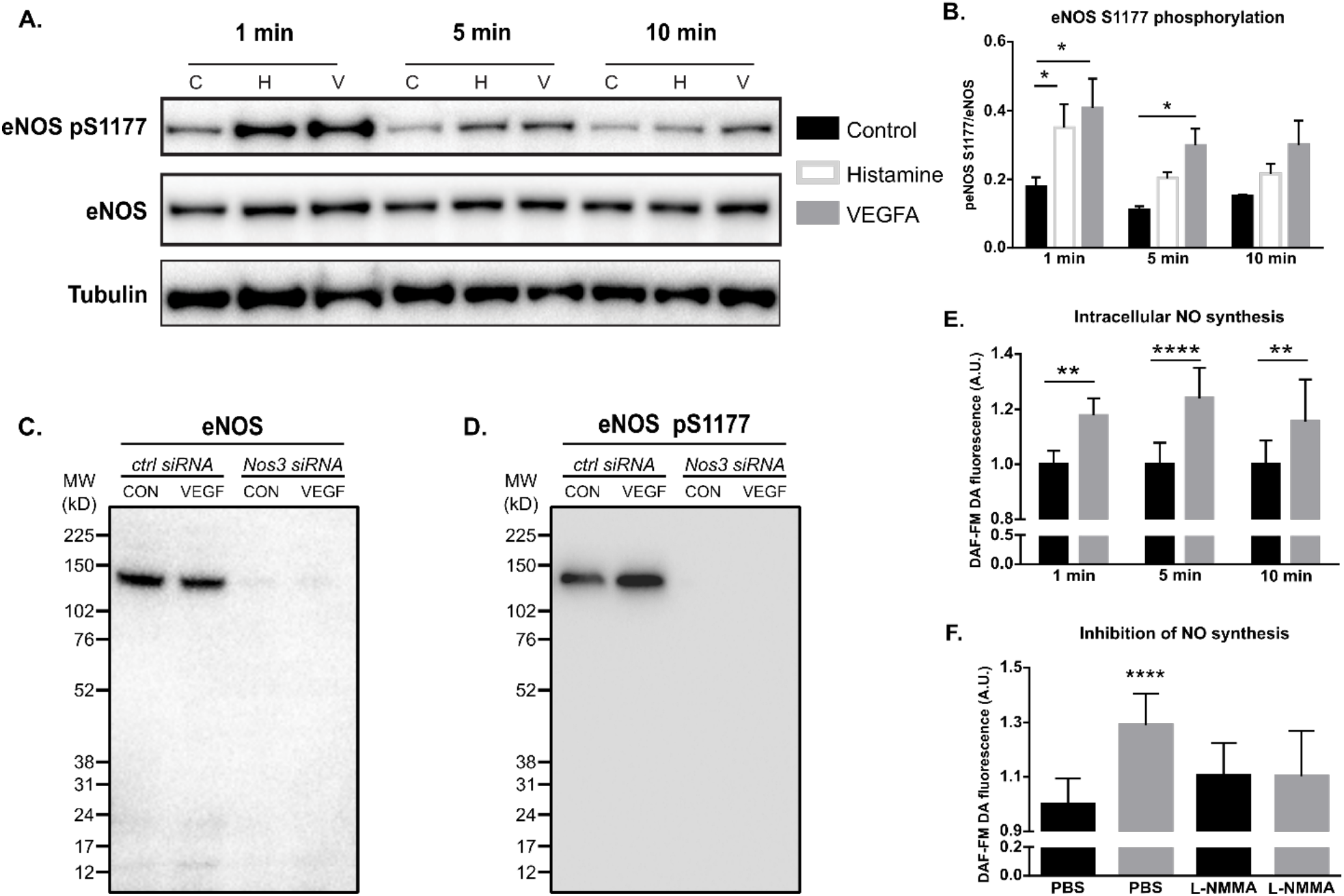
VEGFA induced eNOS phosphorylation and activity *in vitro*. A. Effect of VEGFA (V; 100 ng/mL; 1, 5, 10 min), histamine (H; 10 μM, 1, 5, 10 min) or medium (C, control) on eNOS phosphorylation at S1177 in cultured Human Retinal Microvascular Endothelial Cells (HRMEC). B. Quantification of eNOS pS1177/total eNOS normalized to tubulin. Mean ±S.E.M. n = 3 independent experiments. * = p < 0.05; two-way ANOVA, Sidak’s multiple comparison test. C. Antibody validation by immunoblotting for eNOS on HRMECs transfected with a control siRNA or Nos3-specific siRNA followed by treatment with VEGFA (100 ng/mL; 5 min). D. Antibody validation by immunoblotting for eNOS pS1177 on HRMECs transfected with a control siRNA or Nos3-specific siRNA followed by treatment with VEGFA (100 ng/mL; 5 min). E. Quantification of NO production in HRMECs treated with PBS or VEGFA (100 ng/mL, for 1, 5 or 10 min) using the cell-permeable fluorescent probe DAF-FM DA. F. Quantification of NO production in HRMECs pre-treated with PBS or L-NMMA (1 mM) before VEGFA stimulation (100 ng/mL, 5 min). Mean ±S.E.M. n = 12, 3 independent experiments. *, **, **** = p < 0.05, 0.01, 0.0001; two-way ANOVA, Sidak’s multiple comparison test.

**Figure 2 – supplement figure 2.**
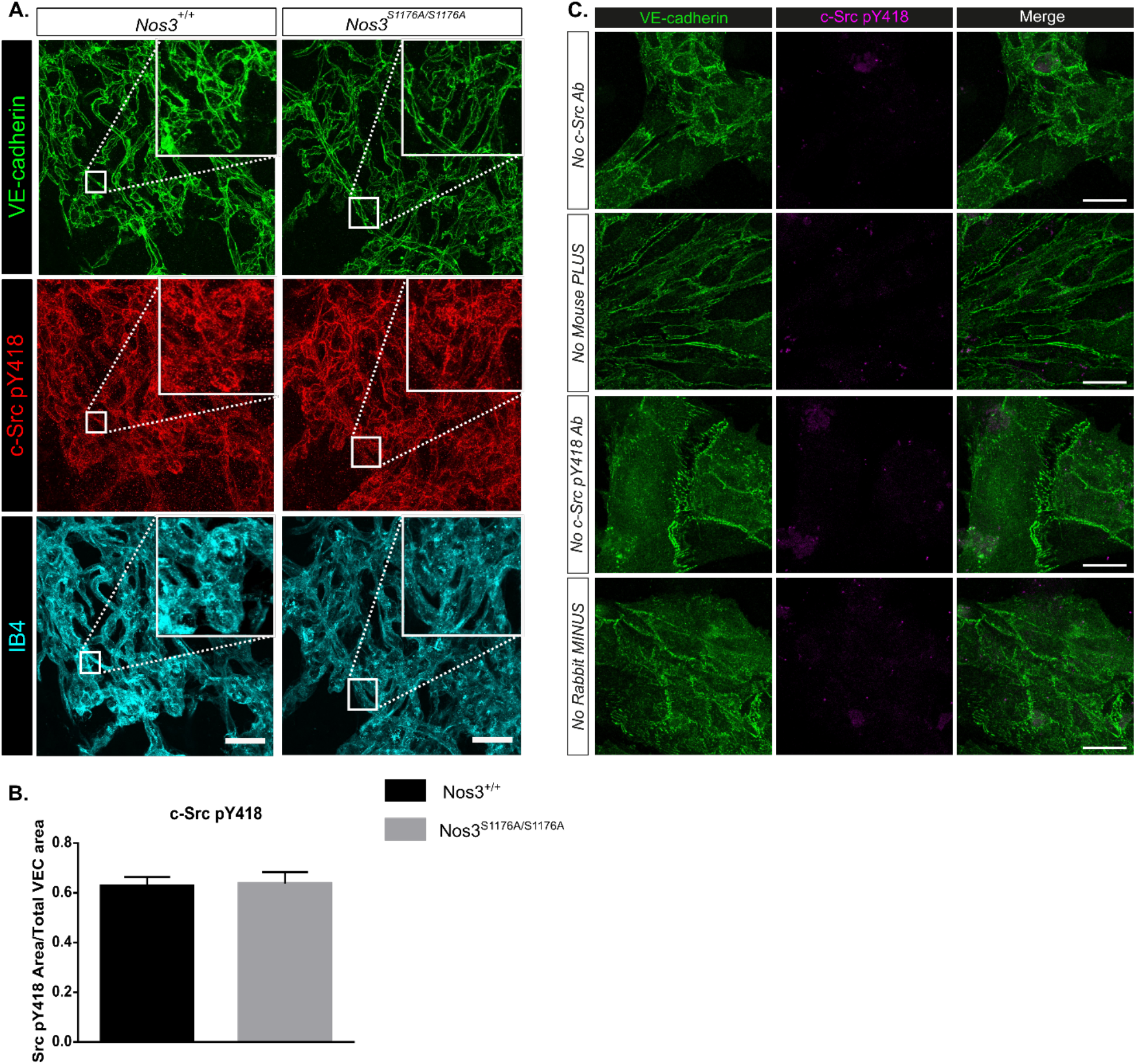
c-Src pY418 immunostaining and PLA controls. A. Representative maximum intensity projections of tufts from Nos3^+/+^ and Nos3^S1176A/S1176A^ retinas immunostained for VE-cadherin (green), c-Src pY418 (red) and IB4 (cyan). B. Quantification of pY418-positive area/ total VE-cadherin area. n = 3 images per group from 3 (*Nos3*^*+/+*^) and 4 (*Nos3*^*S1176A/S1176A*^) mice from 2 separate experiments; t-test. C. Representative images of VE-cadherin staining (green) and proximity ligation controls in isolated mouse lung endothelial cells (iEC) from *Nos3*^*+/+*^ mice. Controls were performed by incubation with only one of the necessary antibodies or, one of the PLUS or MINUS PLA probes (mouse and rabbit secondary antibodies). This allows for the detection of non-specific ligation or uncontrolled rolling circle DNA synthesis. Scale bar = 20 μm.

**Figure 1 – source data 1.**
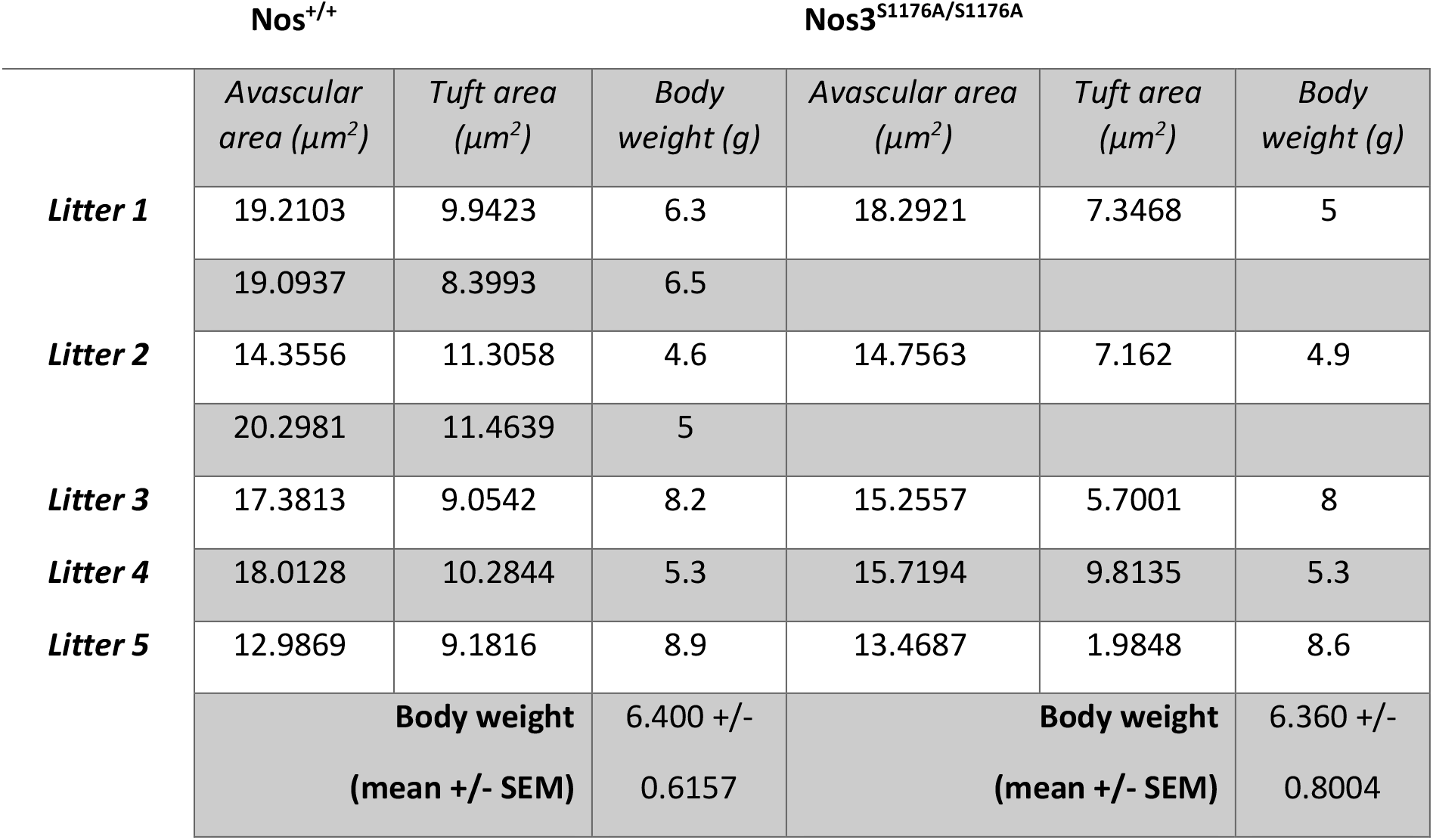
Raw data on retina vascular parameters and body weights from OIR experiments on *Nos*^*+/+*^ and *Nos3*^*S1176A/S1176A*^ mice.

**Figure 3 – source data 1.**
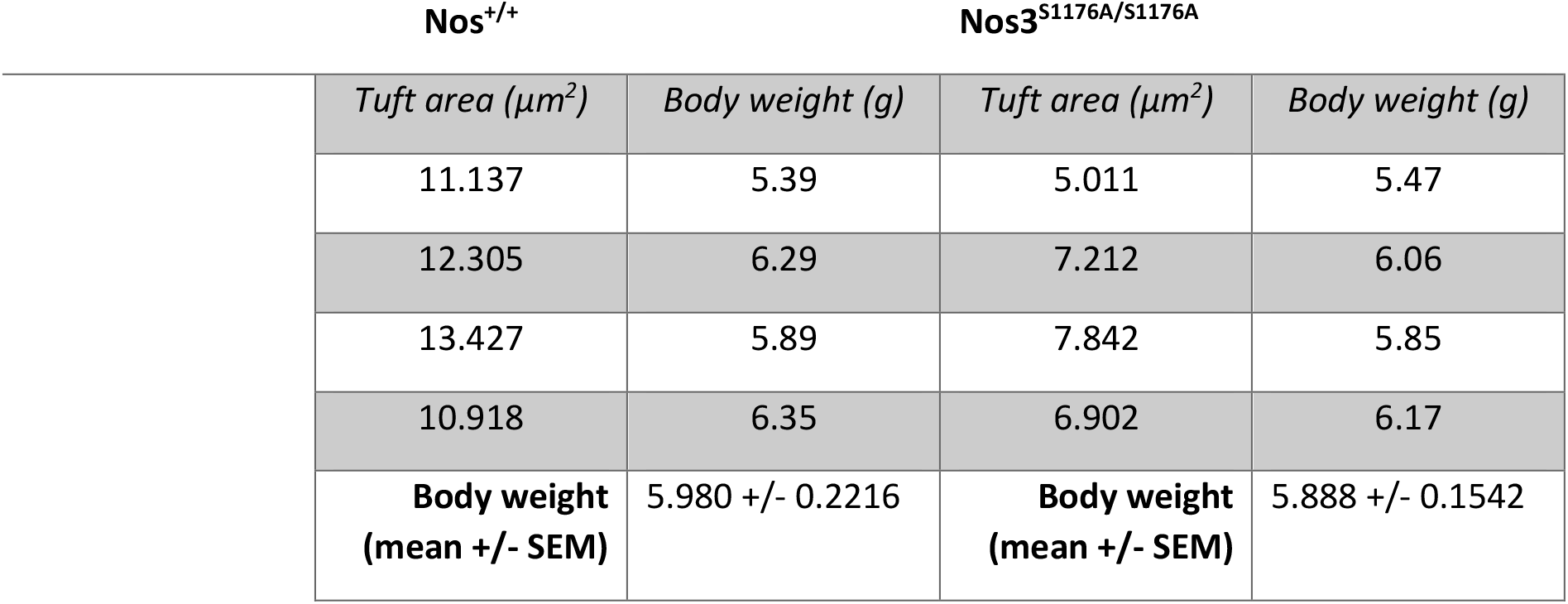
Raw data on retina vascular parameters and body weights from *Nos3*^*+/+*^ and *Nos3*^*S1176A/S1176A*^ mice injected with microspheres.

**Figure 4 – source data 1.**
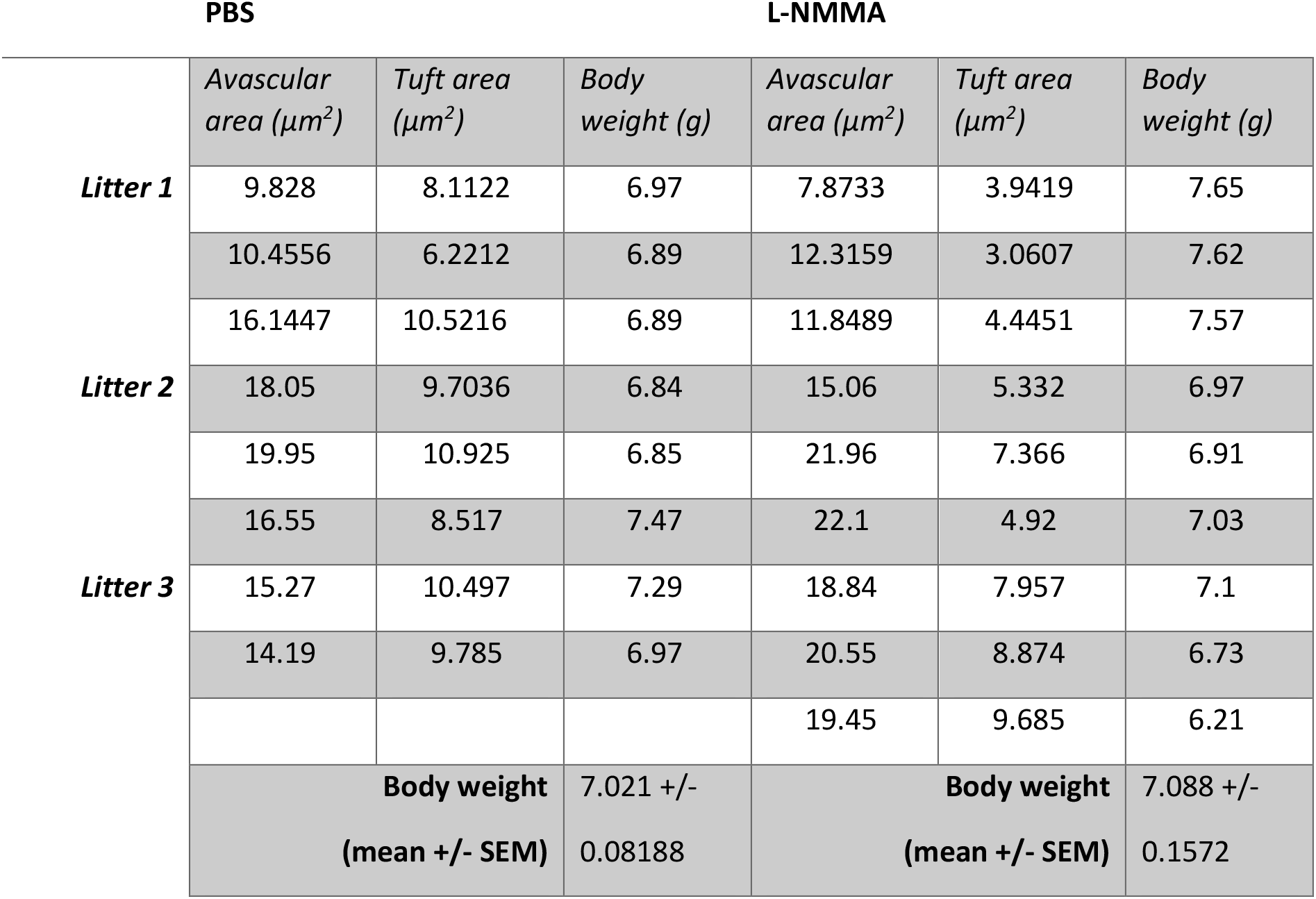
Raw data on retina vascular parameters and body weights from OIR experiments on PBS and L-NMMA treated mice.

**Figure 4 – source data 2.**
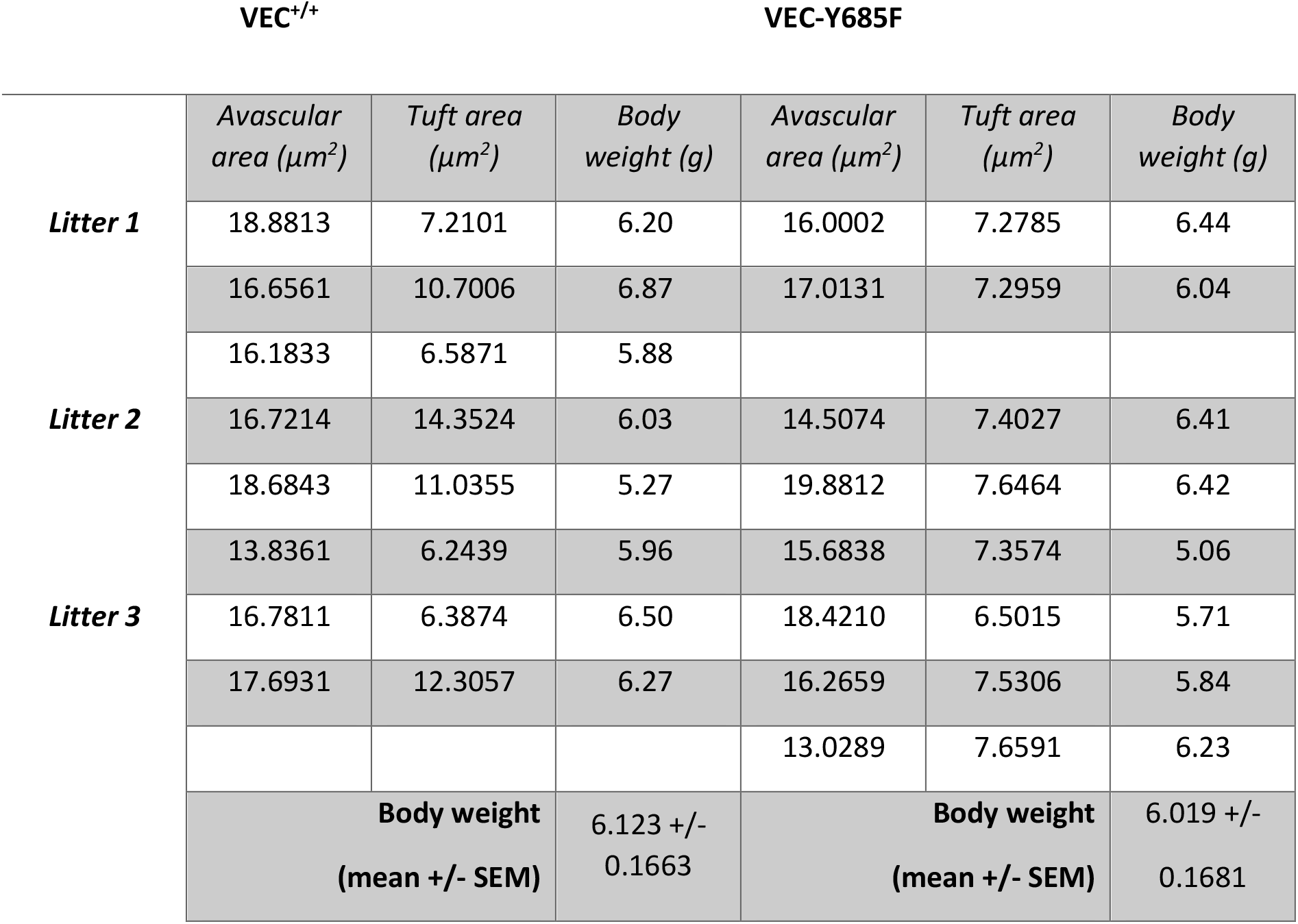
Raw data on retina vascular parameters and body weights fro OIR experiments on VEC^+/+^ and VEC-Y685F mice.

**Figure 5 – source data 1.**
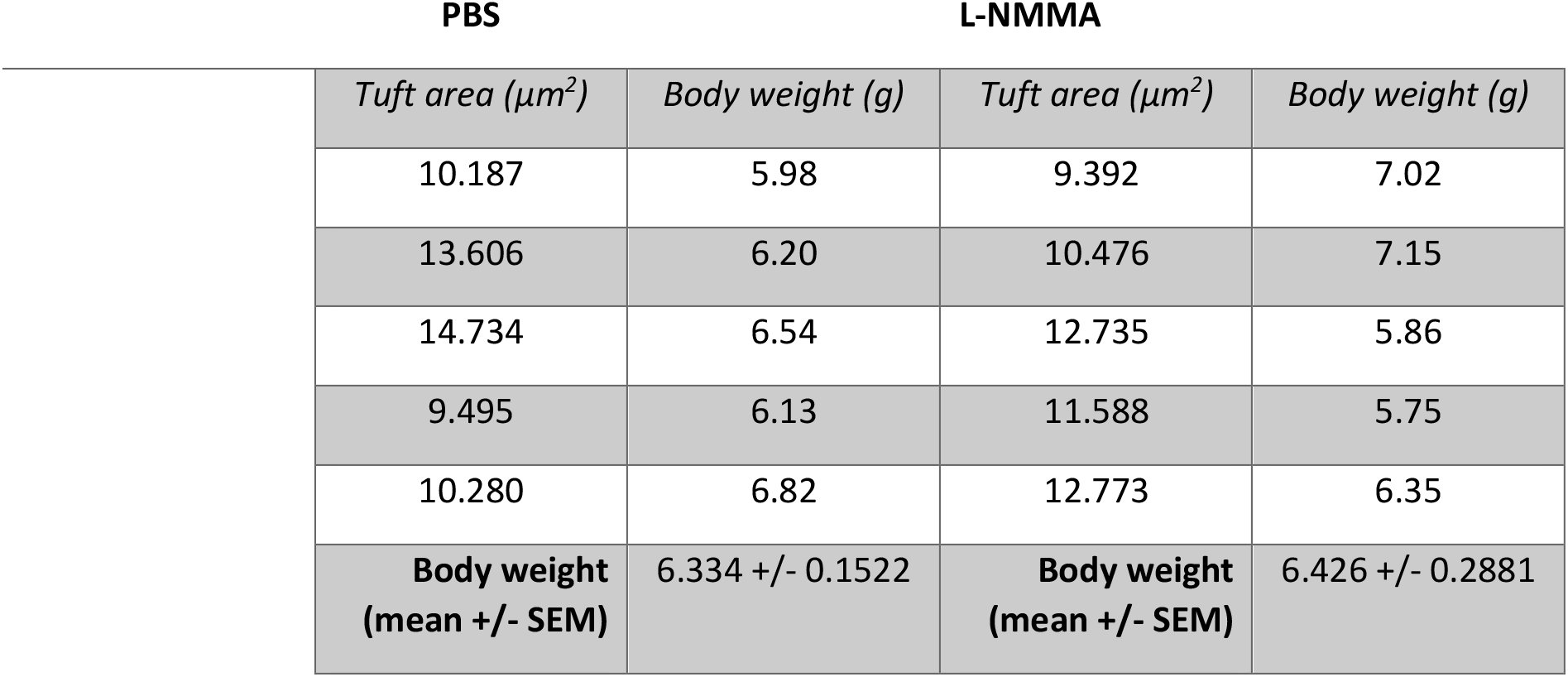
Raw data on retina vascular parameters and body weights from *Nos3*^*+/+*^ mice treated with L-NMMA and injected with microspheres.

